# Mapping Pharmacologically-induced Functional Reorganisation onto the Brain’s Neurotransmitter Landscape

**DOI:** 10.1101/2022.07.12.499688

**Authors:** Andrea I. Luppi, Justine Y. Hansen, Ram Adapa, Robin L. Carhart-Harris, Leor Roseman, Christopher Timmermann, Daniel Golkowski, Andreas Ranft, Rüdiger Ilg, Denis Jordan, Vincent Bonhomme, Audrey Vanhaudenhuyse, Athena Demertzi, Oceane Jaquet, Mohamed Ali Bahri, Naji L.N. Alnagger, Paolo Cardone, Alexander R. D. Peattie, Anne E. Manktelow, Draulio B. de Araujo, Stefano L. Sensi, Adrian M. Owen, Lorina Naci, David K. Menon, Bratislav Misic, Emmanuel A. Stamatakis

## Abstract

To understand how pharmacological interventions can exert their powerful effects on brain function, we need to understand how they engage the brain’s rich neurotransmitter landscape. Here, we bridge microscale molecular chemoarchitecture and pharmacologically-induced macroscale functional reorganisation, by relating the regional distribution of 19 neurotransmitter receptors and transporters obtained from Positron Emission Tomography, and the regional changes in functional MRI connectivity induced by 10 different mind-altering drugs: propofol, sevoflurane, ketamine, LSD, psilocybin, DMT, ayahuasca, MDMA, modafinil, and methylphenidate. Our results reveal that psychoactive drugs exert their effects on brain function by engaging multiple neurotransmitter systems. The effects of both anaesthetics and psychedelics on brain function are organised along hierarchical gradients of brain structure and function. Finally, we show that regional co-susceptibility to pharmacological interventions recapitulates co-susceptibility to disorder-induced structural alterations. Collectively, these results highlight rich statistical patterns relating molecular chemoarchitecture and drug-induced reorganisation of the brain’s functional architecture.

## Introduction

Understanding how the brain orchestrates complex signals across spatial and temporal scales to support cognition and consciousness is a fundamental challenge of contemporary neuroscience. By inducing profound but reversible alterations of brain function, psychoactive compounds provide neuroscientists with the means to manipulate the brain without requiring surgical intervention. In combination with non- invasive brain imaging techniques such as functional MRI, acute pharmacological interventions have therefore emerged as a prominent tool for causal investigation of the relationship between brain and cognitive function in healthy humans^1^.

Mind-altering pharmacological agents also play a fundamental role in modern clinical practice. The invention of anaesthesia was a major milestone in medical history, enabling millions of life-saving surgeries to take place every year^2^. Other drugs that influence the mind without suppressing consciousness, such as the cognitive enhancers modafinil and methylphenidate, have found useful applications in alleviating the cognitive symptoms of syndromes such as ADHD, narcolepsy, and traumatic brain injury (TBI)^3–11^. More recently, classic and “atypical” psychedelics are increasingly being investigated for their potential to provide breakthrough avenues to treat psychiatric conditions, with recent successes in clinical trials heralding a possible end to the current scarcity of therapies for treatment-resistant depression and other neuropsychiatric disorders^12–17^. For these convergent reasons, the effects of anaesthetics, psychedelics, and cognitive enhancers on brain function are becoming the focus of intense investigation, revealing both similarities and differences between them^18–26^.

Pharmacological agents exert their mind-altering effects by tuning the brain’s neurotransmitter landscape. Neurotransmitters engage receptors on neurons’ membrane to mediate the transfer and propagation of signals between cells, modulate the functional configurations of neuronal circuits, and ultimately shape network-wide communication^27–31^. Several psychoactive drugs appear to exert their effects on the mind and brain primarily through one or few specific neurotransmitters: the main action of the general anaesthetic propofol is enhancement of synaptic transmission mediated by GABA-A receptors, a mechanism that is also shared by sevoflurane, which in addition attenuates glutamatergic synaptic signalling (mediated by both AMPA and NMDA receptors)^2, 32–39^. Ketamine (a dissociative anaesthetic at high doses, and atypical psychedelic at low doses) is an NMDA receptor antagonist^40–45;^ the classic psychedelics LSD, psilocybin, and DMT are agonists of the serotonin 2A receptor, with a strong dependence between subjective efficacy and 2A receptor affinity^46–49^.

However, in the words of Sleigh and colleagues, “Linking observed molecular actions for any particular drug with its clinical effects is an abiding pharmacological problem”^50^: knowing the primary molecular target is not sufficient to understand a drug’s effects on brain function, for several reasons. First, given the brain’s intricate, nested feedback loops and recurrent pathways of connectivity, even a relatively selective drug can end up influencing unrelated systems beyond what may be apparent from in vitro studies. Second, most mind-altering compounds are also known to have affinity for other receptors. Indeed, evidence has been accumulating that multiple neurotransmitter influences may be involved in both the neural and subjective experiences induced by many consciousness-altering drugs. In the last year, human neuroimaging studies identified the involvement of the dopaminergic system in both propofol-induced anaesthesia^51^ and the subjective effects of LSD^52^. More broadly, a recent large-scale study, combining receptor expression from transcriptomic data with linguistic processing of several thousand subjective reports of psychedelic use, identified complex multivariate patterns of association between neurotransmitters and their effects on the mind elicited by a wide variety of psychedelics, even for putatively selective agents^53^. At the same time, molecularly different compounds can exert intriguingly similar effects on both the mind and brain: for instance, LSD and (sub-anaesthetic) ketamine can produce subjectively similar effects and changes in terms of structure-function coupling and the complexity of brain activity - despite acting on different pathways^21^. This suggests both divergent and convergent effects of different pharmacological agents on the brain’s rich neurotransmitter landscape.

Finally, the human brain exhibits rich patterns of anatomical, functional, cytoarchitectonic, and molecular variations^54–59^. Such patterns also extend to the regional distribution of different neurotransmitter receptors and transporters, which vary widely not only in terms of their affinity, time-scales, and downstream effects on neuronal excitability, but also their distribution across regions, layers and neuron types^27–30^. Therefore, our knowledge of how a drug influences neurotransmission must take into account the neuroanatomical distribution of its target neurotransmitters - an essential step towards explaining how different neurotransmitters mediate the capacity of different drugs to shape the functional and computational properties of the brain’s architecture^27, 31^.

Here, we sought to address this question in a data-driven way, mapping the neurotransmitter landscape of drug-induced alterations in the brain’s functional connectivity. To do so, we leveraged two unique datasets: (i) a recently assembled collection of in vivo maps of regional receptor expression from 19 different receptors, obtained from PET scanning of over 1200 total subjects, providing the most detailed information about neuromodulators and their spatial distribution available to date^31^; and (ii) resting-state functional MRI (rs-fMRI) data acquired under the effects of the serotonergic psychedelics LSD^60^, psilocybin^61^, DMT (Timmermann et al., under review), ayahuasca^62^, and MDMA^63^; sub-anaesthetic doses of ketamine (acting as an “atypical psychedelic”)^64^ as well as anaesthetic doses (acting as a “dissociative anaesthetic”); the cognitive enhancers modafinil^65^ and methylphenidate^10^; and the anaesthetics sevoflurane^66^ and propofol^67, 68^ (which we compared against pre- anaesthesic baseline as well as post-anaesthetic recovery); representing a total of 382 sessions of pharmacological-MRI from 224 distinct subjects and 10 distinct pharmacological agents. Through pharmacologically modulated rs-fMRI, we can characterise a drug’s effects on the brain’s spontaneous activity, without the interference of any specific task^1^.

Thus, our goal was to obtain a comprehensive mapping between the cortical distributions of neurotransmitters and a set of diverse psychoactive pharmacological agents (covering the range from anaesthetics to psychedelics), in terms of their effects of functional connectivity. There have been other studies looking at the relationships between brain changes induced by one or few psychoactive drugs, and one or few neurotransmitter systems^51, 52, 69–74^ – and a previous effort considering how changes in cerebral blood flow induced by different psychiatric medications depend on the distribution of receptors^75^. However, to our knowledge, this is the largest fMRI study to date not only in terms of the number, variety, and potency of psychoactive pharmacological agents included, but also the breadth and coverage of neurotransmitter systems considered.

## Results

To establish a relationship between neurotransmitter systems and pharmacologically-induced reorganisation of the brain’s functional architecture, we combine two sets of neuroimaging data, each collected from across multiple studies. On one hand, we characterise drug-induced functional reorganisation as the changes in functional connectivity (FC) obtained by contrasting resting-state functional MRI (rs-fMRI) at baseline and under the acute effect of a psychoactive drug. We considered the general anaesthetics propofol (two independent datasets) and sevoflurane; the cognitive enhancers modafinil and methylphenidate; ketamine, acting as both atypical psychedelic (at sub-anaesthetic doses) and as dissociative anaesthetic^24, 75, 76^; and the serotonergic psychedelics lysergic acid diethylamide (LSD), psilocybin, DMT, ayahuasca, and MDMA (Figure 1). For sevoflurane and both propofol datasets, we considered two contrasts: drug versus pre-induction baseline, and drug versus post-anaesthetic recovery (recovery data were not available for ketamine). We followed the same preprocessing and denoising procedure for each dataset, to ensure comparability (see Methods).

**Figure 1.**
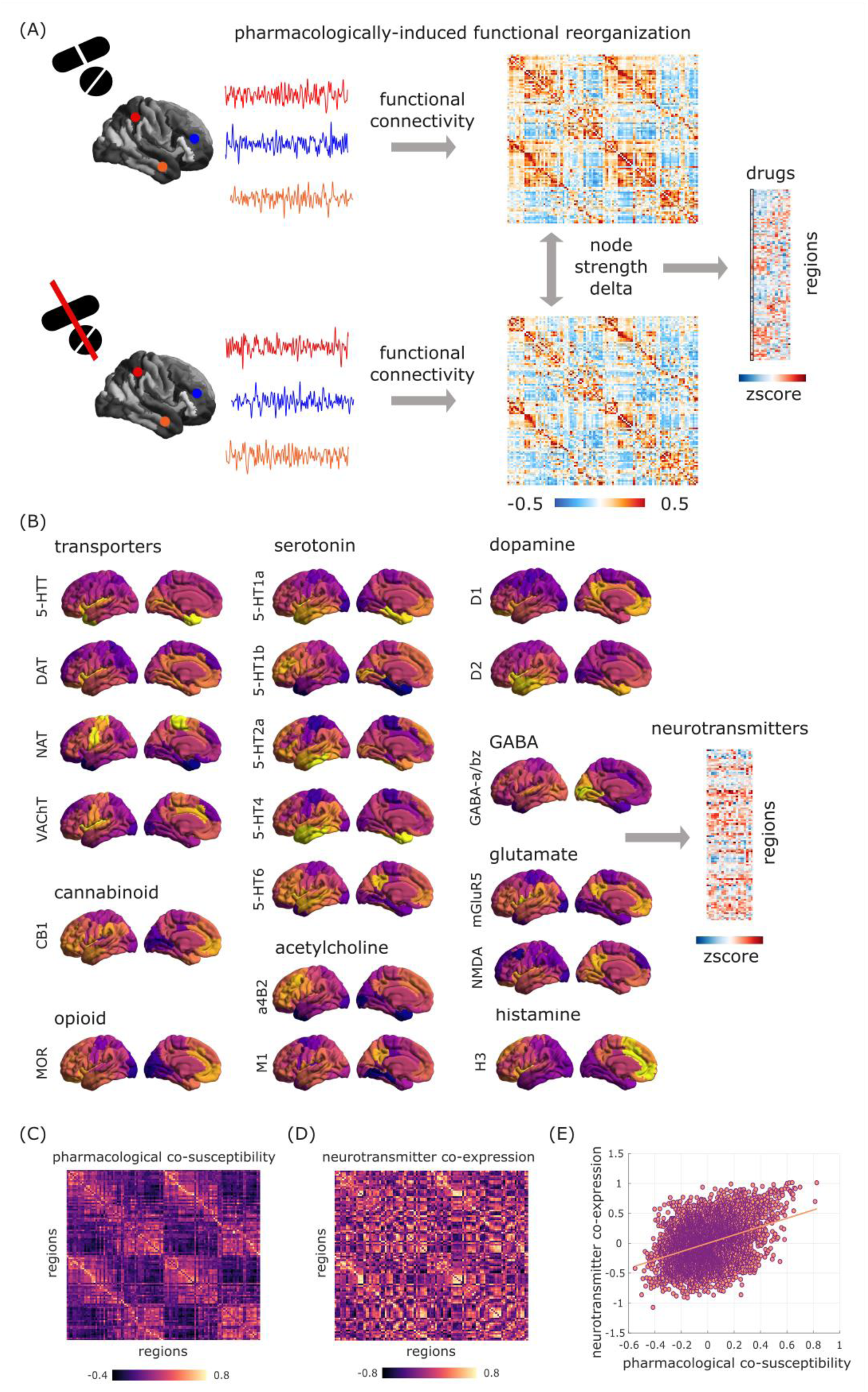
Overview of receptors and pharmacological rs-fMRI data. (A) For each psychoactive drug, its pattern of pharmacologically-induced functional reorganisation is quantified as the average (across subjects) of the within-subject difference in regional FC density between task-free fMRI scans at baseline and under the drug’s effects. The result is a map of 100 cortical regions by 15 drug-related contrasts. (B) Neurotransmitter systems are mapped with Positron Emission Tomography with radioligands for 15 receptors and 4 transporters, resulting in a map of 100 cortical regions by 19 neurotransmitters. Correlating each of these sets of maps against itself yields two region-by-region matrices of pharmacological co-susceptibility (C) and neurotransmitter co-expression (D), respectively, which are significantly correlated even after removing the exponential relationship with Euclidean distance between regions (E).

On the other hand, we consider the cortical distribution of 15 neurotransmitter receptors and 4 transporters, obtained from in vivo Positron Emission Tomography^31^. Overall, 9 neurotransmitter and neuromodulatory systems (“neurotransmitters” for short) are covered: dopamine (D1^77^, D2^78–81^, DAT^82^), norepinephrine (NET^83–86)^, serotonin (5-HT1A^87^, 5-HT1B^87–90,90–92,^ 5-HT2A^93^, 5-HT4^93^, 5-HT6^94, 95^, 5-HTT^93^), acetylcholine (*α*4*β*2^96, 97^, M1^98^, VAChT^99, 100^), glutamate (mGluR5^101, 102^, NMDA^103, 104^), GABA (GABA-A^105^), histamine (H3^106^), cannabinoid (CB1^107–110)^, and opioid (MOR^111^). (Figure 1A,B). Both rs-fMRI and PET maps were parcellated into 100 functionally defined regions according to the Schaefer atlas^112^.

### Brain regions with shared chemoarchitecture also respond similarly across pharmacological perturbations

Receptors and transporters shape the way that neurons respond to neurotransmission and neuromodulatory influences. In turn, psychoactive drugs exert their effects (primarily) by acting on neurotransmitters and neuromodulators. Therefore, we reasoned that everything else being equal, regions that express similar patterns of receptors and transporters should exhibit similar patterns of susceptibility to drug-induced functional reorganisation.

To address this question, we computed matrices of pharmacological co-susceptibility and neurotransmitter co-expression between pairs of regions, by correlating respectively the regional patterns of drug-induced FC changes (across all subjects), and the regional patterns of neurotransmitter expression across all 19 receptor and transporter PET maps. To account for spatial autocorrelation in molecular and FC attributes, we regressed out from both matrices the exponential trend with Euclidean distance^113–116^.

Supporting our hypothesis, we found that pharmacological co-susceptibility is significantly correlated with neurotransmitter profile similarity: the extent to which two regions’ FC patterns are similarly affected by perturbations induced by different psychoactive drugs, is predicted by the extent to which they co-express neurotransmitter receptors and transporters: *rho* = 0.34, *p* < 0.001 after regressing out the effects of Euclidean distance (Figure 1C-E). In other words, regions that exhibit shared chemoarchitecture also respond similarly across pharmacological perturbations.

### Multivariate Receptor-Drug Associations

The previous analysis revealed of a relationship between large-scale patterns of neurotransmitter expression, and large-scale patterns of functional susceptibility to pharmacological perturbations – complementing previous work that identified relationships between individual drugs and individual receptors. However, neither of these two approaches captures the full richness of the two datasets employed here. To obtain a synthesis between these two approaches, we employed a multivariate association technique, Partial Least Squares correlation (PLS, also known as Projection to Latent Structures^117, 118^), which enabled us to identify multivariate patterns of maximum covariance between drug-induced effects on functional connectivity, and the cortical distributions of neurotransmitter expression^119, 120^.

This analysis indicated the presence of two statistically significant latent variables (linear weighted combinations of the original variables) relating pharmacologically- induced functional reorganisation to neurotransmitter profiles, together accounting for nearly 85% of covariance. Significance was assessed against autocorrelation- preserving spin-based null models, embodying the null hypothesis that drug effects and neurotransmitters are spatially correlated with each other purely because of inherent spatial autocorrelation^121–124^ (Figure 2). We further cross-validated this result using a distance-dependent method; out-of-sample *r* = 0.46 for PLS1 and 0.54 for PLS2, both *p* < 0.001 from t-test against spin-based null distributions) (Figure S1).

**Figure 2.**
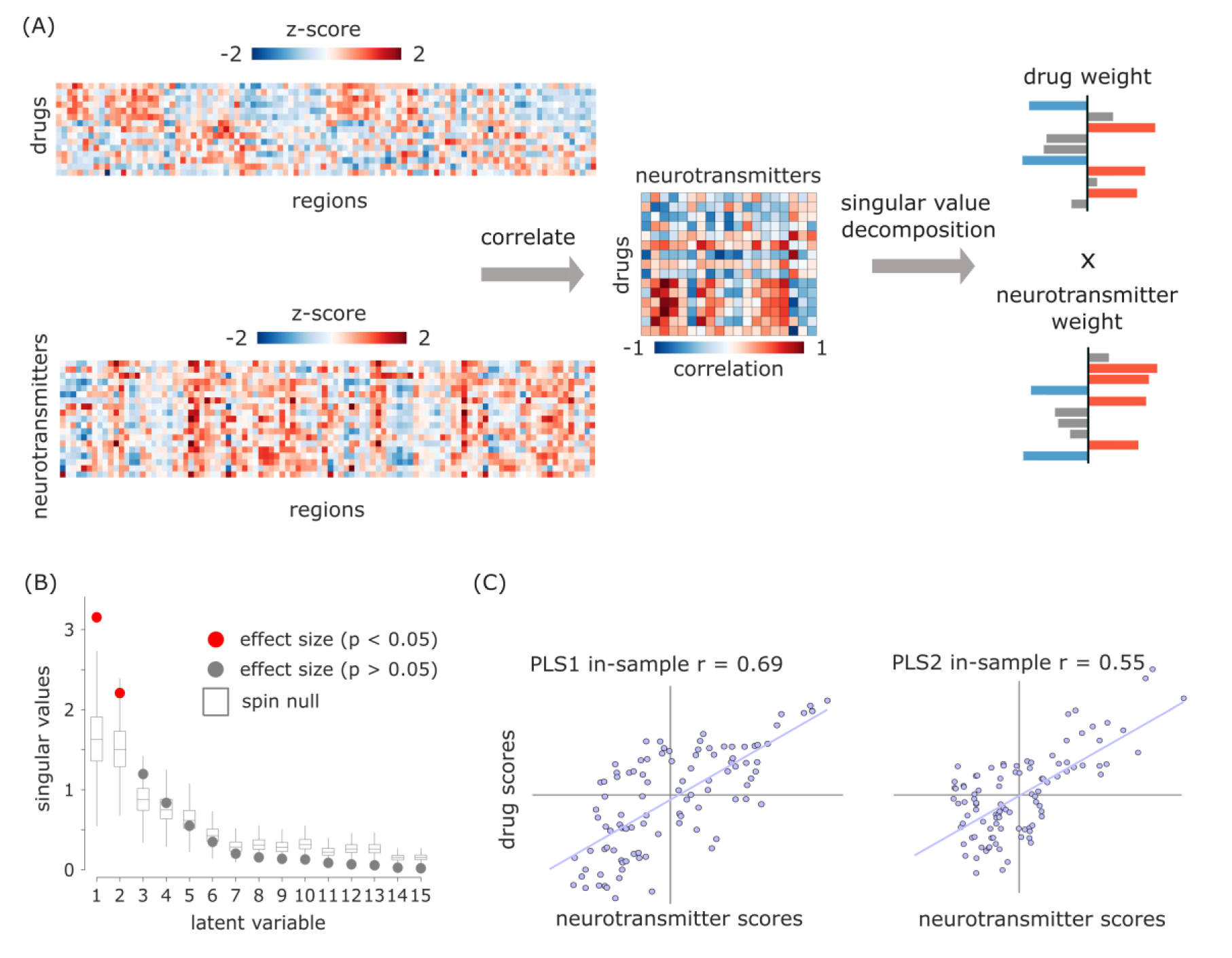
PLS analysis reveals spatially covarying patterns of pharmacologically-induced functional reorganisation and neurotransmitter expression. (A) PLS analysis relates two data domains by correlating the variables across brain regions and subjecting this to singular value decomposition. This results in multiple latent variables: linear weighted combinations of the original variables (neurotransmitter weights and drug weights) that maximally covary with each other. (B) Latent variables are ordered according to effect size (the proportion of covariance explained between neurotransmitter expression and drug-induced functional reorganisation they account for) and shown as red dots. (C) The first two latent variables (PLS1 and PLS2) were statistically significant, with respect to the spatial autocorrelation-preserving null model shown in grey (10,000 permutations). The first latent variable accounted for 57% of covariance, and the second latent variable accounted for 28%. Neurotransmitter (drug) scores are defined as the projection of the original neurotransmitter density (drug-induced FC changes) matrix onto the neurotransmitter (drug) weights, such that each brain region is associated with a neurotransmitter and drug score. By design, neurotransmitter and drug scores correlate highly.

For each latent variable, each brain region is associated with a neurotransmitter and drug score. In turn, neurotransmitter (drug) loadings are defined as the correlation between the PLS-derived score pattern and each neurotransmitter’s density of expression (resp., drug-induced FC changes) across brain regions. Taking into account the first latent variable (PLS1), drug loadings showed a distinction of pharmacological effects into two groups, with all anaesthetics (except ketamine) on one side, and both ketamine datasets dominating the opposite side, together with LSD, ayahuasca, and modafinil (Figure 3A). Neurotransmitter loadings divided the receptors from transporters: at the positive end (orange), the noradrenaline, serotonin and acetylcholine transporters (with the dopamine transporter following closely, but narrowly including zero in its 95% CI); all receptors except NMDA were instead at the negative end (blue), although some included zero in their CI (Figure 3B).

**Figure 3.**
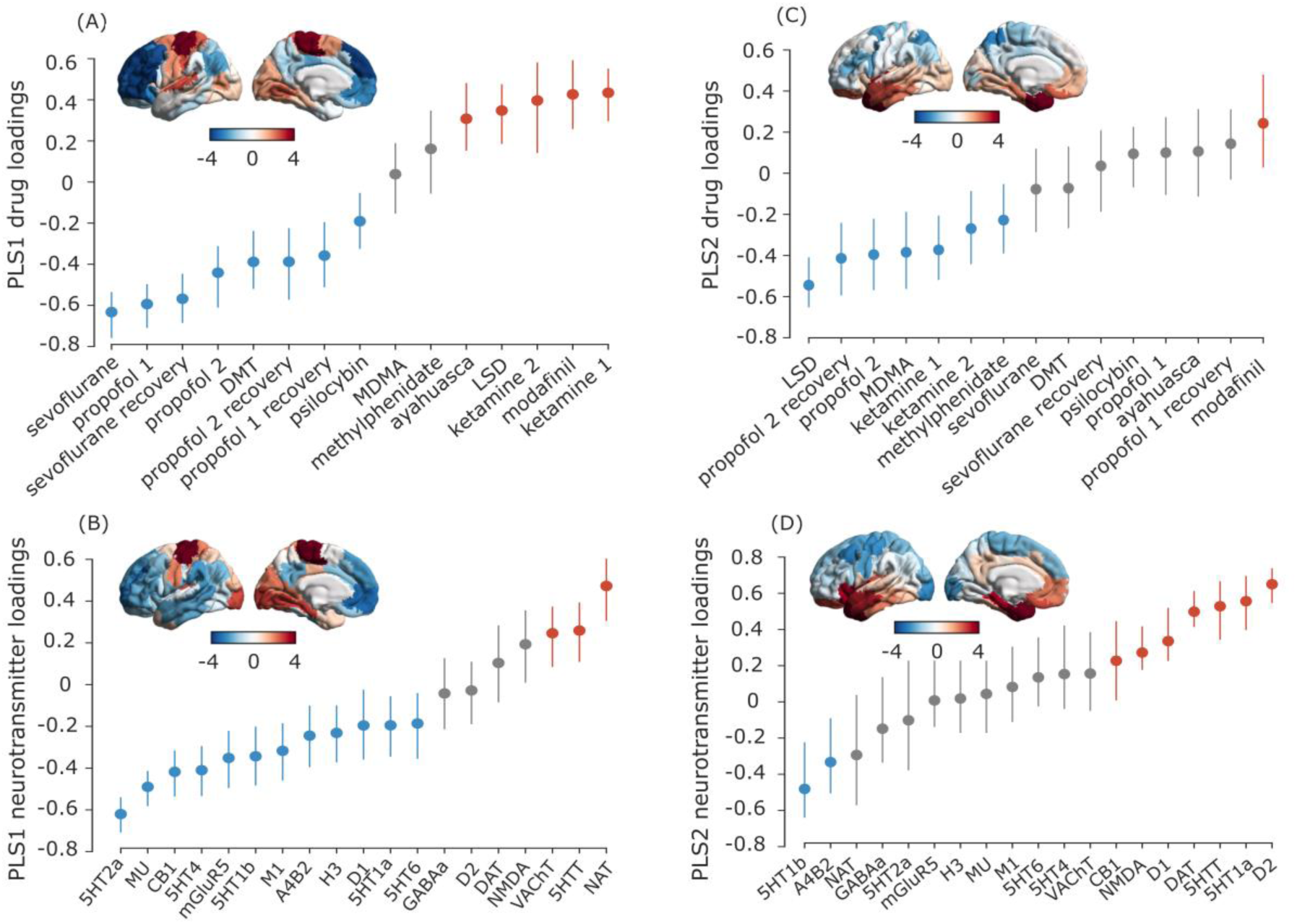
PLS scores and loadings from significant latent variables. (A-B) Scores and loadings for PLS1. (C-D) scores and loadings for PLS2. Brain plots: Drug scores (top row) and neurotransmitter scores (bottom row) for each brain region are obtained by projecting the original neurotransmitter and drug data back onto the PLS analysis-defined drug/neurotransmitter weights, indexing the extent to which a brain region expresses covarying drug/ neurotransmitter patterns. In turn, neurotransmitter (drug) loadings are defined as the Pearson’s correlation between each neurotransmitter’s density of expression (drug-induced FC changes) across brain regions and the PLS analysis-derived score pattern. Error bars indicate 95% confidence interval, and colour indicates direction of the effect: positive (orange), negative (blue), or null (grey). Same-coloured loadings and scores co-vary positively, whereas opposite-coloured drugs and scores co-vary negatively. The label “ketamine 1” refers to the sub-anaesthetic dose, and “ketamine 2” is the anaesthetic dose.

Pertaining to the second latent variable (PLS2), neurotransmitter loadings mainly identified a monoamine-rich end (with dopamine and serotonin), although 5-HT1b occupied the opposite end. However, the drug loadings were less clearly discernible, with modafinil alone at one end, and a mixture of propofol, psychedelics, and both ketamine datasets at the other end. Both neurotransmitter and drug scores markedly separated dorsal and ventral aspects of the brain for this second latent variable (Figure 3).

### Pharmacologically-induced alterations align with functional, anatomical and molecular hierarchies

Neurotransmitter and drug scores (whose spatial similarity PLS is designed to maximise) provide information about the regional distribution of neurotransmitter- drug associations. Neurotransmitters and drugs whose activity correlates positively with the score pattern covary with one another in the positively scored regions, and vice versa for negatively scored regions.

PLS1 scores correspond to the main axis of covariance between neurotransmitter expression and pharmacologically-induced functional reorganisation. For both drug and receptor scores, we observed that their regional distribution reflected the brain’s organisation into intrinsic resting-state networks (RSNs)^125^, setting apart visual and somatomotor cortices from association cortices (Figures 3,4). It is possible that the correspondence of PLS1 scores with RSNs may be in part driven by the fact that these networks are predicated in terms of functional neuroimaging, which we also used to characterise drug-induced functional reorganisation in our data. Therefore, we next sought to determine whether our data-driven topographic patterns reflect other cortical gradients of variation in terms of functional, anatomical, and molecular attributes. To this end, we considered intracortical myelination obtained from T1w/T2w MRI ratio^58^; evolutionary cortical expansion obtained by comparing human and macaque^126^; the principal component of variation in gene expression from the Allen Human Brain Atlas transcriptomic database (“AHBA PC1”)^59, 127^; the principal component of variation in task activation from the NeuroSynth database (“NeuroSynth PC1”)^59, 128^; and the principal gradient of functional connectivity^57^.

**Figure 4.**
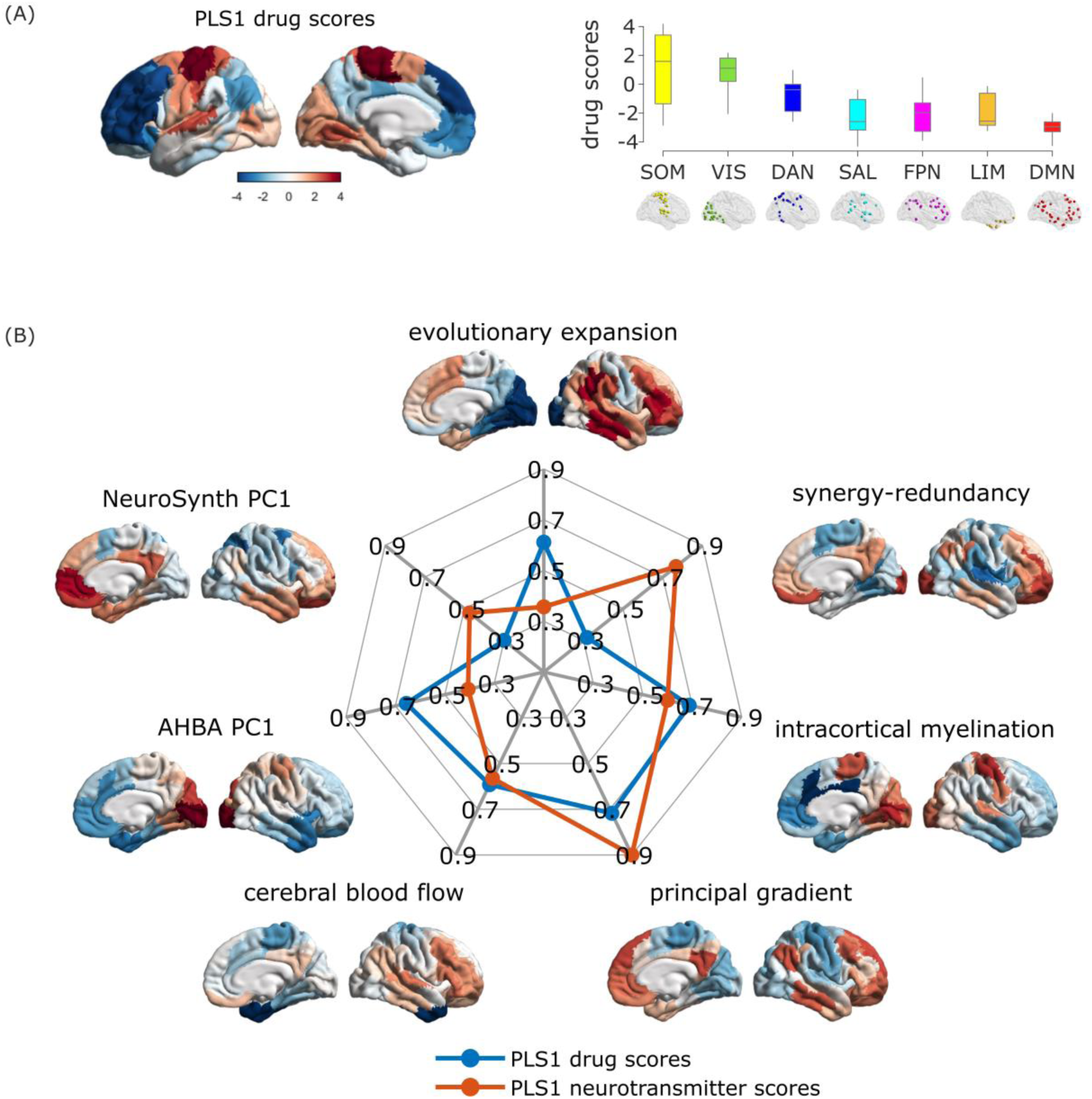
Correspondence between the principal axis of drug-neurotransmitter scores and functional, anatomical and molecular hierarchies. (A) Cortical distribution of drug scores for PLS1, and their association with intrinsic resting-state networks. (B) Radial plot represents the absolute value of the correlation between PLS1 drug and neurotransmiter scores, and each of seven cortical hierarchies obtained from different neuroimaging modalities (note that the myelin and AHBA PC1 maps are reversed with respect to the remaining hierarchies).

Since pharmacological interventions exert their effects on the brain via the bloodstream, we also included a map of cerebral blood flow^54^. Finally, we included a recently derived gradient of regional prevalence of different kinds of information, from redundancy to synergy^129^.

We observed significant correlations (assessed against spin-based null models) between each cortical hierarchy and both neurotransmitter and drug scores for PLS1 (except for PLS1 drug scores versus NeuroSynth PC1; *p_spin_* = 0.054) (Figure 4). The scores for PLS2 instead identified a ventral-dorsal pattern of regional variation (Figure 3 and Figure S2), which did not significantly correlate with any of the canonical gradients of hierarchical organisation (all *p >* 0.05 against spin-based null models, except for PLS2 drug scores versus NeuroSynth PC1; *p_spin_* = 0.013).

### Neurotransmitter landscape of pharmacologically-induced functional reorganisation

Taking into account the first two PLS latent variables shows how each drug-specific pattern of pharmacologically-induced functional reorganisation can be interpreted in terms of contributions from different receptors (note that sign is arbitrary) (Figure 5). As already shown in Figure 3, the first latent variable revealed a stark division between transporters and receptors, which discriminates between traditional anaesthetics and other psychoactive substances. In terms of pharmacological alterations, the non-monoaminergic end of the second latent variable loaded onto drugs with relatively stronger effects on subjective experiences (the higher doses of anaesthetic, including ketamine, and LSD and MDMA). However, methylphenidate and sub-anaesthetic ketamine also loaded onto this end of the second latent variable. Altogether, we find that the first latent variable captures a strong relationship between drug interventions and receptor systems that is both biologically relevant and aligns with the functional organisation of the brain.

**Figure 5.**
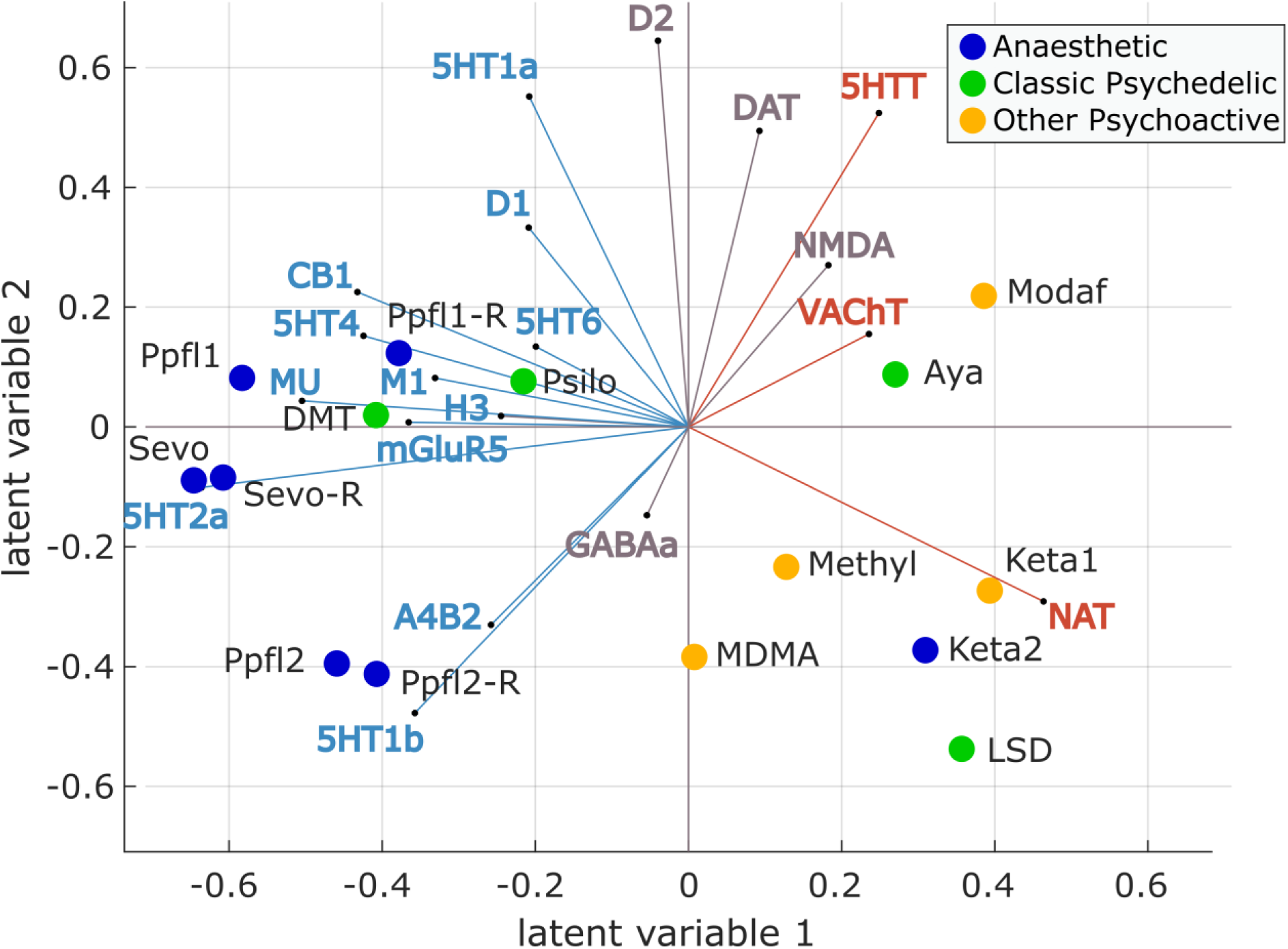
Biplot of neurotransmitters and pharmacological agents. Each drug is represented as a point reflecting its projection onto the first two latent variables of the PLS analysis, color-coded based on its effects on subjective experience (anaesthetic, psychedelic, or other psychoactive). Each neurotransmitter receptor and transporter is represented as a vector in the same 2D space, color-coded by loading onto PLS1 as shown in Figure 3 (orange for positive; blue for negative; and grey if the 95% CI intersects zero). For both propofol (Ppfl) and ketamine (keta), the number refers to the dataset, with 1 identifying the weaker dose, and 2 identifying the stronger dose. Figure S3 shows the drugs and neurotransmitters separately.

### Co-susceptibility to pharmacological and pathological alterations

Finally, we wondered if the functional co-susceptibility of different regions to transient pharmacological perturbations may provide a functional proxy for their co-susceptibility to structural perturbations resulting from different neurological, neurodevelopmental, and psychiatric disorders. To this end, we combined 11 spatial maps of cortical thickness abnormalities made available by the Enhancing Neuro Imaging Genetics Through Meta Analysis (ENIGMA) consortium^130^: 22q11.2 deletion syndrome^131^, attention-deficit/hyperactivity disorder^132^, autism spectrum disorder^133^, idiopathic generalized epilepsy^134^, right temporal lobe epilepsy^134^, left temporal lobe epilepsy^134^, depression^135^, obsessive-compulsive disorder^136^, schizophrenia^137^, bipolar disorder^138^, and Parkinson’s disease^139^. For simplicity, we refer to diseases, disorders, and conditions as “disorders” throughout the text. The cortical abnormality maps summarise contrasts between over 21,000 adult patients and 26,000 controls, collected following identical processing protocols to ensure maximal comparability^130^.

Following the same procedure used to obtain the region x region matrices of pharmacological co-susceptibility and neurotransmitter co-expression in Figure 1, we obtained a region x region matrix of co-susceptibility to disorder-induced cortical abnormality by correlating the regional patterns of cortical abnormality across all 11 disorders^113^ (Figure 6A,B). Correlating this matrix of regional co-susceptibility to disease-associated perturbations against the previously derived matrix of regional co-susceptibility to pharmacological perturbations, we found a statistically significant relationship (Spearman’s *rho* = 0.31, *p* < 0.001 after regressing out the effect of Euclidean distance) (Figure 6C). This result goes beyond a recent demonstration that molecular similarity and disorder similarity are correlated^113^, by showing that a correlation also exists between different kinds of perturbations: anatomical and pharmacological.

**Figure 6.**
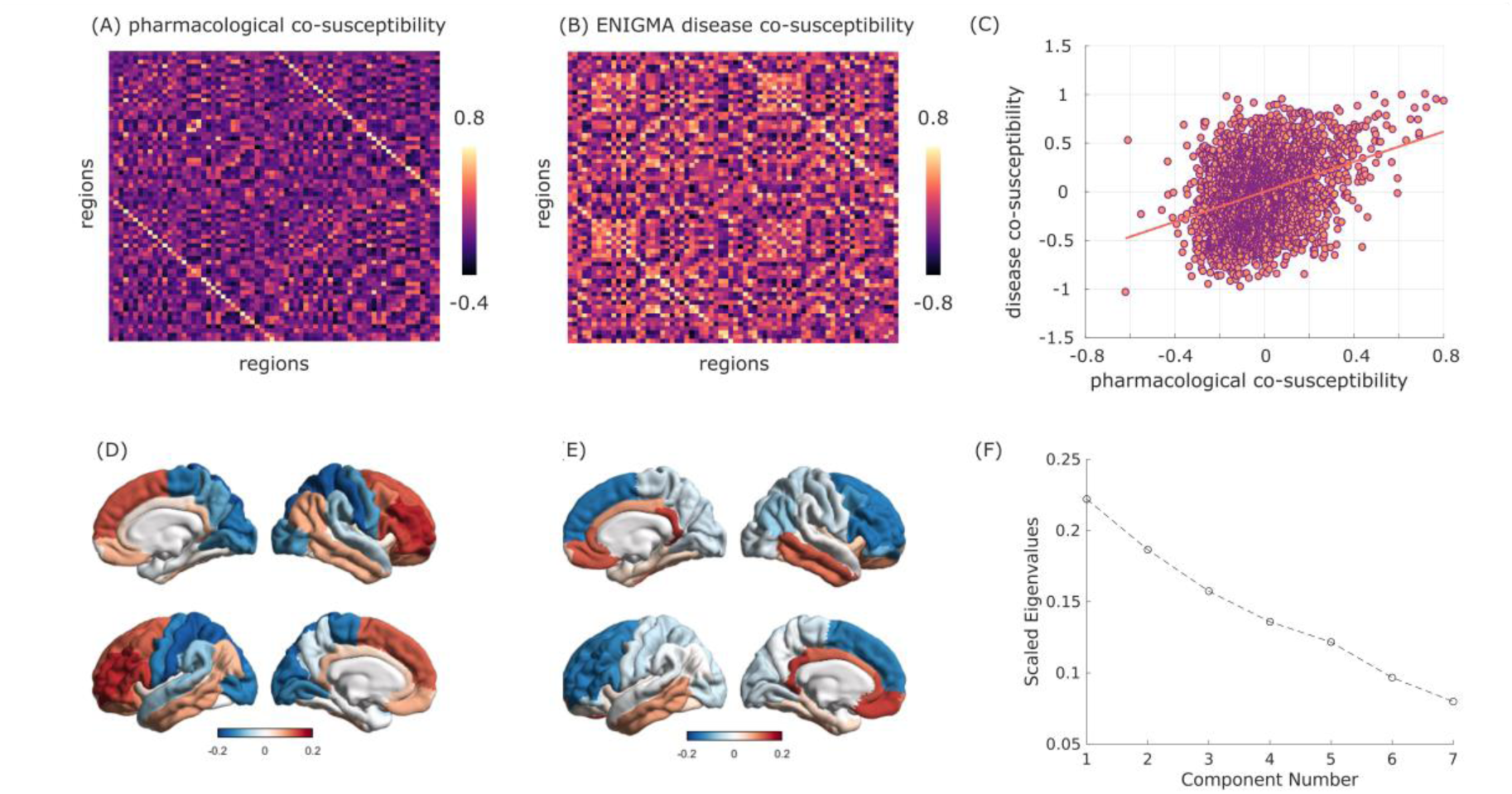
Co-susceptibility to pharmacological and pathological alterations. Brain regions that are similarly affected by pharmacology, in terms of functional reorganisation (A) are also similarly affected across disorders (B), in terms of cortical thickness abnormalities. This relationship persists after regressing out the exponential trend with Euclidean distance (C). (D-E) First two principal gradients of regional joint susceptibility to pharmacological and neuropsychiatric and neurological alterations, obtained from diffusion map embedding. (F) Scree plot of the scaled eigenvalues from diffusion map embedding versus number of components.

This observation suggests that there may be common patterns of regional co-susceptibility to perturbations, whether structural or functional. To explore this possibility explicitly, we resorted to a non-linear dimensionality reduction algorithm, diffusion map embedding^57, 140^, to obtain joint gradients of variation from pharmacological and disease-associated co-susceptibility using a recently developed method for network fusion^141^. We found that the main axis of variation in regional joint susceptibility to pharmacological and neuropsychiatric alterations corresponds to the well-known principal gradient of functional connectivity^57^, setting apart unimodal from transmodal cortices, reminiscent of the PLS1 scores (Figure 6D). The second gradient instead sets apart anterior dorsal and more ventral parts of the brain, reminiscent of the PLS2 scores (Figure 6E). Together, these two gradients account for nearly half of the variation in regional co-susceptibility (Figure 6F).

When applying diffusion map embedding to the matrix of pharmacological co-susceptibility only, we found that the first two gradients of variation in regional pharmacological susceptibility coincide with the two principal gradients of functional connectivity of Margulies *et al*^57^ (Figure S4): the first gradient sets apart unimodal from transmodal cortices, coinciding with the first gradient of joint susceptibility, whereas the second gradient is anchored in visual cortex at one end, and somatomotor cortex at the other end. This observation suggests that co-susceptibility to pharmacological perturbations recapitulates intrinsic functional architecture, as well as the co-susceptibility to disorder-induced structural perturbations.

### Validation and additional analyses

Results for a different cortical parcellation (Lausanne-114^142^) are provided as Supplementary Information (Figures S5, S6); likewise, we show a qualitatively similar mapping between drugs and neurotransmitters on the first two latent variables, even when the methylphenidate dataset (obtained from patients rather than healthy controls) is excluded (Figure S7).

We also performed a separate PLS analysis on the subcortex, which revealed a monoamine- (dopamine- and serotonin-) dominated principal latent variable, positively associated with the drugs having strongest effects, and separating caudate, putamen, and anterior thalamus from posterior thalamus, hippocampus and amygdala (Figure S8).

## Discussion

Here, we characterised how mind-altering pharmacological agents engage the brain’s rich neurotransmitter landscape to exert their effects on brain function. We mapped the functional chemoarchitecture of the human brain, by developing a computational framework to relate the regional reorganisation of fMRI functional connectivity induced by 10 different mind-altering drugs, and the cortical distribution of 19 neurotransmitter receptors and transporters obtained from PET^31^. This approach allowed us to discover large-scale spatial gradients relating pharmacologically-induced changes in functional connectivity to the underlying neurotransmitter systems. By relating microscale molecular chemoarchitecture and macroscale functional reorganisation induced by drugs with potent acute effects on the mind, our results provide a first step to bridge molecular mechanisms and their effects on subjective experience, cognition, and behaviour, via their effects on the brain’s functional architecture.

Using our computational framework, we found that psychoactive drugs are best understood in terms of contributions from multiple neurotransmitter systems. We also found that anaesthetics and psychedelics/cognitive enhancers are largely opposite in terms of their association with neurotransmitters in the cortex – although not without exceptions. Remarkably, the effects of mind-altering drugs are topographically organised along multiple hierarchical gradients of brain function, anatomy, and neurobiology. Finally, we found that co-susceptibility to pharmacological perturbations recapitulates co-susceptibility to disorder-induced structural perturbations.

The diverse mapping between drug-induced functional reorganisation and neurotransmitters that we recovered (Figure 5) clearly shows the power of our multivariate approach to detect both expected and novel relationships between drugs and neurotransmitters. Many of the drugs considered here are known to have varied molecular targets, beyond the primary ones through which they exert their effects. The present results add another dimension to recent work employing a similar multivariate approach to relate gene expression of receptors with subjective reports of psychedelic experiences, which also found widespread involvement of multiple receptors^53^. In addition, the drugs we considered here have profound effects on the mind after a single acute dose^143^, from cognitive enhancement to hallucinations to the suppression of consciousness altogether. Such far-reaching effects are accompanied by sometimes drastic repercussions on brain function and dynamics: it stands to reason that such widespread reorganisation would not leave many neurotransmitter pathways unaffected - even those that are not directly involved in generating the altered state in question.

The broadly opposite characterisation of traditional anaesthetics and most psychedelics is aligned with their respective effects on the complexity of brain activity and connectivity, which is reduced by GABA-ergic anaesthesia but increased by LSD, ayahuasca and psychedelic doses of ketamine, as well as other psychedelics^18, 21, 23, 62, 144–154^. Similarly, psychedelics (including sub-anaesthetic ketamine) and anaesthetics were recently shown to exert opposite effects on structure-function coupling: whereas anaesthesia increases the dependence of brain activity on the underlying structural network, LSD, psilocybin, and sub-anaesthetic ketamine induce fMRI BOLD signals that are increasingly liberal with respect to the underlying structural network organisation^21^. Intriguingly, we found that anaesthetic doses of ketamine align more closely with psychedelic (sub-anaesthetic) doses of ketamine than with anaesthetics such as propofol and sevoflurane. Although the ketamine-anaesthetised volunteers were behaviourally unresponsive, as for the traditional anaesthetics, they subsequently reported a wide range of vivid hallucinatory experiences^75^. Thus, as per ketamine’s characterisation as a dissociative anaesthetic, in subjective terms their consciousness was not suppressed but rather profoundly altered in a manner more similar to psychedelics than anaesthetics. Therefore, the neurotransmitter signatures of the two levels of ketamine align with the molecular effects and subjective effects, rather than with the behavioural effects.

The main division we observed in terms of neurotransmitters is between receptors and transporters, which displayed opposite associations with drug-induced effects. Specifically pertaining to PLS1, we found that transporters covary with cognitive enhancers and most psychedelics in primary sensory and motor regions, whereas receptors covary with GABA-ergic anaesthetics in transmodal association cortices. Hierarchical organisation of pharmacologically-induced functional reorganisation stands to reason based on prior evidence: both psychedelics and anaesthetics have been shown to have potent effects on the activity and connectivity of higher-order association cortices, and the default mode network in particular^18, 60, 61, 66, 75, 155–157^. In addition, serotonergic psychedelics also exert powerful influences on the visual cortex at the other end of the cortical hierarchy^60^, and as a result they have been shown to induce a “flattening” of the principal gradient of functional connectivity^158^.

Having established that the effects of mind-altering drugs are hierarchically organised, the question then becomes: why should mind-altering drugs exert their effects in such a hierarchically organised fashion? Multiple aspects of neuroanatomy may contribute to this effect. First, the principal component of variation of receptor expression is itself organised along the brain’s sensory-to-association hierarchical axis^27^ - and so is, for instance, the distribution of the serotonin 2A receptor, the main direct target of serotonergic psychedelics^31^. Second, transmodal cortices are characterised by increased excitability^159^ and a predominance of feedback efferent connections^27^: combined with their high diversity of receptor expression across layers^27^, these regions may be especially susceptible to receive and amplify multiple pharmacological influences.

Third, we observed that for the first latent variable, neurotransmitter and drug scores also correlate with the map of regional cerebral blood flow; since ultimately the bloodstream is how drugs reach their regional molecular targets, greater cerebral blood flow in transmodal cortices may facilitate especially high availability of the drug in these regions (although it should be noted that some drugs can also have effects on heart rate and neuro-vascular coupling). Finally, regions of transmodal cortex have high neuron and synapse density^129, 160^ and tend to have numerous, far-reaching, and diversely distributed anatomical connections^161^, as well as the highest prevalence of synergistic (complementary) interactions with the rest of the brain^129^, so that any effects that are exerted in these regions may quickly reverberate throughout the whole cortex.

To summarise, we conjecture that the hierarchical organisation of pharmacologically-induced changes in FC may be explained as follows: transmodal association cortices are especially diverse in their receptor profiles, and rich in some key receptors; in addition to being more susceptible to pharmacological intervention due to higher expression of receptors, blood flow is poised to bring greater amounts of drug to these very cortices; and once these cortices’ activity is perturbed, the perturbation can reverberate widely, thanks to their widespread and diverse connectivity. Of course, the drugs we included were chosen precisely because of their powerful effects on cognition and subjective experience, so it stands to reason that their effects should align with the division between primary and higher-order cortices (which also aligns with the principal component of variation obtained from NeuroSynth term-based meta-analysis). In other words, drugs whose effects on functional connectivity are less selective for higher versus lower ends of the cortical hierarchy may simply be less likely to exert mind-altering effects of the kind that we chose to focus on in this work.

More broadly, we found that pairs of regions that are more similar in terms of their susceptibility to pharmacologically-induced FC changes, are also more similar in their susceptibility to cortical alterations associated with a variety of neuropsychiatric disorders. This observation suggests a broader pattern of both pharmacological (acute) and neuroanatomical (chronic) susceptibility across regions. We speculate that this joint susceptibility may be related to regional relevance for cognitive function: indeed, we found that this joint vulnerability can be understood in terms of two multimodal principal gradients of variation over the cortex: one of them resembling the principal gradient of functional connectivity (and principal latent variable of neurotransmitter-drug association), and the other anchored in dorsal prefrontal cortex at one end, and temporal cortex at the other. The association between disorder co-susceptibility and co-susceptibility to pharmacologically-induced functional reorganisation sheds new light on recent evidence that the principal gradient of neurotransmitter expression is particularly relevant for predicting a wide spectrum of disease-specific cortical morphology^113^, by showing that this observation extends to the effects of engaging different receptors. This interpretation is further supported by our own evidence that pharmacological perturbations are shaped by neurotransmitter co-expression.

The results reported here open new possibilities for data-driven, multivariate mapping between the brain’s high-dimensional neurotransmitter landscape and the effects of potent pharmacological interventions on the brain’s functional architecture. Crucially, neuropsychiatric disorders and candidate pharmacological treatments for them ultimately need to exert their effects on cognition and behaviour by influencing brain function. In this light, it is intriguing that susceptibility to disorder-related cortical abnormalities correlates with susceptibility to pharmacological intervention. This observation suggests that regions that are structurally most vulnerable to disease (which presumably in turn shapes their functional architecture) may also be the ones that are most susceptible to re-balancing of their functional organisation by an appropriate choice of pharmacological intervention. This work represents the necessary first step towards identifying novel and perhaps unexpected associations between drugs and neurotransmitters, as well as elucidating the known ones in a data-driven manner.

### Limitations and future directions

Although the main strength of our study is our extensive coverage of both neurotransmitters and pharmacological data, it is important to acknowledge that neither is complete: in particular, our sample did by no means exhaustively include all mind-altering drugs that have been studied: prominent additions for future work may include the psychedelic kappa opioid receptor agonist salvinorin-A^162^, the sedative dexmedetomidine, an alpha-2 receptor agonist^163–165^ – but also alcohol or caffeine, arguably the two most widely used psychoactive substances. We also acknowledge that the pharmacological datasets included here come from limited samples that have been studied before, and future replication in different datasets with the same drugs (as we have done here for propofol) would also be desirable.

The datasets included here come from different sources and locations, and were acquired under a variety of conditions. We endeavoured to mitigate scanner and acquisition differences by re-preprocessing all data with the same pipeline, and following uniform denoising procedures, rather than following the various pipelines originally employed by each group. Further mitigation of the acquisition differences between datasets should come from our within-subject design in healthy individuals (except for the methylphenidate dataset, which comes from TBI patients^10^; although we showed that our results remain qualitatively the same if this dataset is excluded; Fig. S6). Nevertheless, we cannot exclude some residual influence of such differences on our results (e.g., eyes open versus closed; the ayahuasca data were acquired at a lower field strength of 1.5T; the TRs varied from 1.671s for modafinil, to 3s for psilocybin). Similar considerations about the differences between datasets apply for the PET data, as discussed in detail in the original publication collecting the PET maps^31^. Likewise, the coverage of neurotransmitter receptors and transporters, though the most extensive available to date and obtained *in vivo* rather than post-mortem, is far from exhaustive. The same limitation also applies to the ENIGMA disorder data^130^: many more disorders, diseases, and conditions exist than the ones considered here. And although the ENIGMA consortium provides datasets from large samples with standardised pipelines, ensuring robust results, the patient populations may exhibit co-morbidities and/or be undergoing treatment. In addition, the available maps do not directly reflect changes in tissue volume, but rather the effect size of patient-control statistical comparisons, in terms of only one low resolution cortical-only parcellation.

In addition to the inevitable limitations of analysing large-scale datasets from multiple sites, there are also limitations of our analytic framework. Although we report a macroscale spatial association between neurotransmitter expression and pharmacologically-induced functional reorganisation that is statistically unexpected based on autocorrelation alone, caution is warranted when drawing inferences from statistical results to the underlying biology. We used linear models that assume independence between observations – an assumption that mostly does not hold in the brain, given the possibility of nonlinear effects in how drugs exert their effects on the brain’s intricately connected neurotransmitter systems. To mitigate this limitation, throughout this work we triangulated towards a robust statistical mapping between neurotransmitters and drugs by combining cross-validation and conservative null models that account for the spatial dependencies between regions^59^.

Another limitation is that, due to data availability and well documented differences in PET radioligand uptake between cortical and subcortical structures^31, 166, 167^, our work was mainly restricted to the cortex, and we considered the subcortex separately. The thalamus, brainstem, and other subcortical structures are prominently involved in mediating cortico-cortical interactions and the effects of both psychedelics, anaesthetics, and cognitive enhancers^16, 18, 51, 66, 163, 168–173^. We expect that future work suitable data for whole-brain coverage (ideally including cerebellum and brainstem) may provide richer insights than the sum of their individual contributions.

More broadly, the other main limitation of this work is its correlational nature: receptors and drugs were mapped in separate cohorts of individuals, and identifying spatially correlated patterns does not guarantee the causal involvement of the neurotransmitters in question. Experimental interventions will be required to conclusively demonstrate causal involvement, and elucidate the underlying neurobiological pathways. However, we emphasise that our results generate empirically testable hypotheses about which neurotransmitters may be involved with the macroscale effects of different drugs on brain function. Such hypotheses may be tested experimentally, but also *in silico*: whole-brain computational modelling is becoming increasingly prominent as a tool to investigate the causal mechanisms that drive brain activity and organisation in healthy and pathological conditions ^174–177^. Crucially, the more biologically-inspired models (e.g. dynamic mean-field) can also be enriched with further information, such as regional myelination ^159^, or the regional distribution of specific receptors and ion channels obtained from PET or transcriptomics ^71–73, 178, 179^, to reflect neurotransmitter influences. This approach may complement experimental manipulations, making it possible to systematically evaluate the causal effects of combinations of different neuromodulators on the brain’s functional connectivity.

## Conclusion

Here, we mapped the functional chemoarchitecture of the human brain, by relating the regional changes in fMRI functional connectivity induced by 10 different mind-altering drugs, and the regional distribution of 19 neurotransmitter receptors and transporters obtained from PET. This work provides a computational framework to characterise how mind-altering pharmacological agents engage the brain’s rich neurotransmitter landscape to exert their effects on brain function. Our analytic workflow could find application across the breadth of human cognitive and clinical neuroscience, with the potential to shed light on alterations of neurotransmission underlying neuropsychiatric conditions, which are known to involve a combination of anatomical and neurochemical imbalances. More broadly, our framework could also find fruitful application for data-driven prediction of the effects of candidate drugs on the brain: the mapping between neurotransmitters and pharmacological effects on brain function offers an indispensable biological lens that can reveal neurotransmitter targets for therapeutic intervention. In summary, we demonstrate that diverse patterns of neurotransmitter expression are variously engaged by an array of potent pharmacological interventions, ultimately manifesting as a large-scale hierarchical axis. Collectively, these results highlight a statistical link between molecular dynamics and drug-induced reorganisation of functional architecture.

## Materials and Methods

### Description of datasets

#### Propofol

Propofol (2,6-diisopropylphenol) is perhaps the most common agent used for intravenous induction and maintenance of general anaesthesia^32^. One of the chief reasons for its widespread use, both in the operating room and for scientific studies, is propofol’s rapid action, which allow for precise titration and therefore greater control over the induction and emergence process. Additionally, propofol has minimal effects on both regional cerebral blood flow^180^, and the coupling between blood flow and metabolism^181^, thereby reducing the number of potential confounding effects. Propofol is a potent agonist of inhibitory GABA-A receptors, directly activating them as well as increasing their affinity for agonists^33, 34^, leading to suppressed neuronal activity. Propofol also blocks Na+ channels, inhibiting glutamate release^182^ and more broadly it inhibits neurotransmitter release at presynaptic terminals^183^. There is also some evidence that it may affect the dopaminergic system^51, 184, 185^. Here, we included two independent propofol datasets.

##### Western University dataset: Recruitment

The Western University (“Western”) propofol data were collected between May and November 2014 at the Robarts Research Institute, Western University, London, Ontario (Canada), and have been published before^18, 151, 186, 187^. The study received ethical approval from the Health Sciences Research Ethics Board and Psychology Research Ethics Board of Western University (Ontario, Canada). Healthy volunteers (n=19) were recruited (18–40 years; 13 males). Volunteers were right-handed, native English speakers, and had no history of neurological disorders. In accordance with relevant ethical guidelines, each volunteer provided written informed consent, and received monetary compensation for their time. Due to equipment malfunction or physiological impediments to anaesthesia in the scanner, data from n=3 participants (1 male) were excluded from analyses, leaving a total n=16 for analysis^18^.

##### Western University dataset: Study protocol

Resting-state fMRI data were acquired at different propofol levels: no sedation (Awake), Deep anaesthesia (corresponding to Ramsay score of 5) and also during post-anaesthetic recovery. As previously reported^18^, for each condition fMRI acquisition began after two anaesthesiologists and one anaesthesia nurse independently assessed Ramsay level in the scanning room. The anaesthesiologists and the anaesthesia nurse could not be blinded to experimental condition, since part of their role involved determining the participants’ level of anaesthesia. Note that the Ramsay score is designed for critical care patients, and therefore participants did not receive a score during the Awake condition before propofol administration: rather, they were required to be fully awake, alert and communicating appropriately. To provide a further, independent evaluation of participants’ level of responsiveness, they were asked to perform two tasks: a test of verbal memory recall, and a computer-based auditory target-detection task. Wakefulness was also monitored using an infrared camera placed inside the scanner.

Propofol was administered intravenously using an AS50 auto syringe infusion pump (Baxter Healthcare, Singapore); an effect-site/plasma steering algorithm combined with the computer-controlled infusion pump was used to achieve step-wise sedation increments, followed by manual adjustments as required to reach the desired target concentrations of propofol according to the TIVA Trainer (European Society for Intravenous Aneaesthesia, eurosiva.eu) pharmacokinetic simulation program. This software also specified the blood concentrations of propofol, following the Marsh 3- compartment model, which were used as targets for the pharmacokinetic model providing target-controlled infusion. After an initial propofol target effect-site concentration of 0.6 *µ*g mL^-1^, concentration was gradually increased by increments of 0.3 *µ*g mL^1^, and Ramsay score was assessed after each increment: a further increment occurred if the Ramsay score was lower than 5. The mean estimated effect-site and plasma propofol concentrations were kept stable by the pharmacokinetic model delivered via the TIVA Trainer infusion pump. Ramsay level 5 was achieved when participants stopped responding to verbal commands, were unable to engage in conversation, and were rousable only to physical stimulation. Once both anaesthesiologists and the anaesthesia nurse all agreed that Ramsay sedation level 5 had been reached, and participants stopped responding to both tasks, data acquisition was initiated. The mean estimated effect-site propofol concentration was 2.48 (1.82- 3.14) *µ*g mL^-1^, and the mean estimated plasma propofol concentration was 2.68 (1.92- 3.44) *µ*g mL^-1^. Mean total mass of propofol administered was 486.58 (373.30- 599.86) mg. These values of variability are typical for the pharmacokinetics and pharmacodynamics of propofol. Oxygen was titrated to maintain SpO2 above 96%.

At Ramsay 5 level, participants remained capable of spontaneous cardiovascular function and ventilation. However, the sedation procedure did not take place in a hospital setting; therefore, intubation during scanning could not be used to ensure airway security during scanning. Consequently, although two anaesthesiologists closely monitored each participant, scanner time was minimised to ensure return to normal breathing following deep sedation. No state changes or movement were noted during the deep sedation scanning for any of the participants included in the study^18^. Propofol was discontinued following the deep anaesthesia scan, and participants reached level 2 of the Ramsey scale approximately 11 minutes afterwards, as indicated by clear and rapid responses to verbal commands. This corresponds to the “recovery” period.

As previously reported^18^, once in the scanner participants were instructed to relax with closed eyes, without falling asleep. Resting-state functional MRI in the absence of any tasks was acquired for 8 minutes for each participant. A further scan was also acquired during auditory presentation of a plot-driven story through headphones (5- minute long). Participants were instructed to listen while keeping their eyes closed. The present analysis focuses on the resting-state data only; the story scan data have been published separately^188^ and will not be discussed further here.

##### Western University dataset: MRI Data Acquisition

As previously reported^18^, MRI scanning was performed using a 3-Tesla Siemens Tim Trio scanner (32-channel coil), and 256 functional volumes (echo-planar images, EPI) were collected from each participant, with the following parameters: slices = 33, with 25% inter-slice gap; resolution = 3mm isotropic; TR = 2000ms; TE = 30ms; flip angle = 75 degrees; matrix size = 64x64. The order of acquisition was interleaved, bottom-up. Anatomical scanning was also performed, acquiring a high-resolution T1- weighted volume (32-channel coil, 1mm isotropic voxel size) with a 3D MPRAGE sequence, using the following parameters: TA = 5min, TE = 4.25ms, 240x256 matrix size, 9 degrees flip angle^18^.

##### Cambridge University dataset: Recruitment

The Cambridge University (“Cambridge”) propofol dataset is described in detail in a previous publication^67^. Sixteen healthy volunteer subjects were initially recruited for scanning. In addition to the original 16 volunteers, data were acquired for nine additional participants using the same procedures, bringing the total number of participants in this dataset to 25 (11 males, 14 females; mean age 34.7 years, SD = 9.0 years). Ethical approval for these studies was obtained from the Cambridgeshire 2 Regional Ethics Committee, and all subjects gave informed consent to participate in the study. Volunteers were informed of the risks of propofol administration, such as loss of consciousness, respiratory and cardiovascular depression. They were also informed about more minor effects of propofol such as pain on injection, sedation and amnesia. In addition, standard information about intravenous cannulation, blood sampling and MRI scanning was provided.

##### Cambridge University dataset: Study protocol

Three target plasma levels of propofol were used - no drug (Awake), 0.6 mg/ml (Mild sedation) and 1.2 mg/ml (Moderate sedation). Scanning (rs-fMRI) was acquired at each stage, and also at Recovery; anatomical images were also acquired. The level of sedation was assessed verbally immediately before and after each of the scanning runs. Propofol was administered intravenously as a “target controlled infusion” (plasma concentration mode), using an Alaris PK infusion pump (Carefusion, Basingstoke, UK). A period of 10 min was allowed for equilibration of plasma and effect-site propofol concentrations. Blood samples were drawn towards the end of each titration period and before the plasma target was altered, to assess plasma propofol levels. In total, 6 blood samples were drawn during the study. The mean (SD) measured plasma propofol concentration was 304.8 (141.1) ng/ml during mild sedation, 723.3 (320.5) ng/ml during moderate sedation and 275.8 (75.42) ng/ml during recovery. Mean (SD) total mass of propofol administered was 210.15 (33.17) mg, equivalent to 3.0 (0.47) mg/kg. Two senior anaesthetists were present during scanning sessions and observed the subjects throughout the study from the MRI control room and on a video link that showed the subject in the scanner. Electrocardiography and pulse oximetry were performed continuously, and measurements of heart rate, non-invasive blood pressure, and oxygen saturation were recorded at regular intervals.

##### Cambridge University dataset: MRI Data Acquisition

The acquisition procedures are described in detail in the original study^37^. Briefly, MRI data were acquired on a Siemens Trio 3T scanner (WBIC, Cambridge). For each level of sedation, 150 rs-fMRI volumes (5 min scanning) were acquired. Each functional BOLD volume consisted of 32 interleaved, descending, oblique axial slices, 3 mm thick with interslice gap of 0.75 mm and in-plane resolution of 3 mm, field of view = 192x192 mm, repetition time = 2000 ms, acquisition time = 2 s, time echo = 30 ms, and flip angle 78. T1-weighted structural images at 1 mm isotropic resolution were also acquired in the sagittal plane, using an MPRAGE sequence with TR = 2250 ms, TI = 900 ms, TE = 2.99 ms and flip angle = 9 degrees, for localization purposes. During scanning, volunteers were instructed to close their eyes and think about nothing in particular throughout the acquisition of the resting state BOLD data. Of the 25 healthy subjects, 15 were ultimately retained (7 males, 8 females): 10 were excluded, either because of missing scans (n=2), or due of excessive motion in the scanner (n=8, 5mm maximum motion threshold). For the analyses presented in this paper, we only considered the Awake, Moderate (i.e., loss of behavioural responsiveness) and Recovery resting-state scans.

#### Sevoflurane

Sevoflurane is an inhalational anaesthetic: specifically, a halogenated ether. Although its exact molecular mechanisms of action are yet to be fully elucidated, in vivo and in vitro evidence indicates that it acts primarily via GABA-A receptors^2, 35–38^, but also interacts with NMDA, AMPA^39, 189, 190^ and nicotinic ACh receptors^191, 192^. Additionally, electrophysiologic investigation suggests possible affinity for as well as Na+, K+ and hyperpolarization-activated cyclic nucleotide-gated (HCN) channels^193^.

##### Sevoflurane dataset: Recruitment

The data included here have been published before ^66, 194–196^, and we refer the reader to the original publication for details^66^. The ethics committee of the medical school of the Technische Universität München (München, Germany) approved the current study, which was conducted in accordance with the Declaration of Helsinki. Written informed consent was obtained from volunteers at least 48 h before the study session. Twenty healthy adult men (20 to 36 years of age; mean, 26 years) were recruited through campus notices and personal contact, and compensated for their participation in the study.

Before inclusion in the study, detailed information was provided about the protocol and risks, and medical history was reviewed to assess any previous neurologic or psychiatric disorder. A focused physical examination was performed, and a resting electrocardiogram was recorded. Further exclusion criteria were the following: physical status other than American Society of Anesthesiologists physical status I, chronic intake of medication or drugs, hardness of hearing or deafness, absence of fluency in German, known or suspected disposition to malignant hyperthermia, acute hepatic porphyria, history of halothane hepatitis, obesity with a body mass index more than 30 kg/m2, gastrointestinal disorders with a disposition for gastroesophageal regurgitation, known or suspected difficult airway, and presence of metal implants. Data acquisition took place between June and December 2013.

##### Sevoflurane dataset: Study protocol

Sevoflurane concentrations were chosen so that subjects tolerated artificial ventilation (reached at 2.0 vol%) and that burst-suppression (BS) was reached in all participants (around 4.4 vol%). To make group comparisons feasible, an intermediate concentration of 3.0 vol% was also used. In the MRI scanner, volunteers were in a resting state with eyes closed for 700s. Since EEG data were simultaneously acquired during MRI scanning ^66^ (though they are not analysed in the present study), visual online inspection of the EEG was used to verify that participants did not fall asleep during the pre-anaesthesia baseline scan. Sevoflurane mixed with oxygen was administered via a tight-fitting facemask using an fMRI-compatible anaesthesia machine (Fabius Tiro, Dräger, Germany). Standard American Society of Anesthesiologists monitoring was performed: concentrations of sevoflurane, oxygen and carbon dioxide, were monitored using a cardiorespiratory monitor (DatexaS/3, General electric, USA). After administering an end-tidal sevoflurane concentration (etSev) of 0.4 vol% for 5 min, sevoflurane concentration was increased in a stepwise fashion by 0.2 vol% every 3 min until the participant became unconscious, as judged by the loss of responsiveness (LOR) to the repeatedly spoken command “squeeze my hand” two consecutive times. Sevoflurane concentration was then increased to reach an end-tidal concentration of approximately 3 vol%. When clinically indicated, ventilation was managed by the physician and a laryngeal mask suitable for fMRI (I-gel, Intersurgical, United Kingdom) was inserted. The fraction of inspired oxygen was then set at 0.8, and mechanical ventilation was adjusted to maintain end-tidal carbon dioxide at steady concentrations of 33 ± 1.71 mmHg during BS, 34 ± 1.12 mmHg during 3 vol%, and 33 ± 1.49 mmHg during 2 vol% (throughout this article, mean ± SD). Norepinephrine was given by continuous infusion (0.1 ± 0.01 µg ꞏkg^−1^ ꞏ min^−1^) through an intravenous catheter in a vein on the dorsum of the hand, to maintain the mean arterial blood pressure close to baseline values (baseline, 96 ± 9.36 mmHg; BS, 88 ± 7.55 mmHg; 3 vol%, 88 ± 8.4 mmHg; 2 vol%, 89 ± 9.37 mmHg; follow-up, 98 ± 9.41 mmHg). After insertion of the laryngeal mask airway, sevoflurane concentration was gradually increased until the EEG showed burst-suppression with suppression periods of at least 1,000 ms and about 50% suppression of electrical activity (reached at 4.34 ± 0.22 vol%), which is characteristic of deep anaesthesia. At that point, another 700s of electroencephalogram and fMRI was recorded. Further 700s of data were acquired at steady end-tidal sevoflurane concentrations of 3 and 2 vol%, respectively, each after an equilibration time of 15 min. In a final step, etSev was reduced to two times the concentration at LOR. However, most of the subjects moved or did not tolerate the laryngeal mask any more under this condition: therefore, this stage was not included in the analysis^66^.

Sevoflurane administration was then terminated, and the scanner table was slid out of the MRI scanner to monitor post-anaesthetic recovery. The volunteer was manually ventilated until spontaneous ventilation returned. The laryngeal mask was removed as soon as the patient opened his mouth on command. The physician regularly asked the volunteer to squeeze their hand: recovery of responsiveness was noted to occur as soon as the command was followed. Fifteen minutes after the time of recovery of responsiveness, the Brice interview was administered to assess for awareness during sevoflurane exposure; the interview was repeated on the phone the next day. After a total of 45 min of recovery time, another resting-state combined fMRI-EEG scan was acquired (with eyes closed, as for the baseline scan). When participants were alert, oriented, cooperative, and physiologically stable, they were taken home by a family member or a friend appointed in advance.

##### Sevoflurane dataset: MRI Data Acquisition

Although the original study acquired both functional MRI (fMRI) and electroencephalographic (EEG) data, in the present work we only considered the fMRI data. Data acquisition was carried out on a 3-Tesla magnetic resonance imaging scanner (Achieva Quasar Dual 3.0T 16CH, The Netherlands) with an eight-channel, phased-array head coil. The data were collected using a gradient echo planar imaging sequence (echo time = 30 ms, repetition time (TR) = 1.838 s, flip angle = 75°, field of view = 220 × 220 mm^2^, matrix = 72 × 72, 32 slices, slice thickness = 3 mm, and 1 mm interslice gap; 700-s acquisition time, resulting in 350 functional volumes). The anatomical scan was acquired before the functional scan using a T1-weighted MPRAGE sequence with 240 × 240 × 170 voxels (1×1×1 mm voxel size) covering the whole brain. A total of 16 volunteers completed the full protocol and were included in our analyses; one subject was excluded due to high motion, leaving N=15 for analysis. Here, we used fMRI data from the Awake, 3% vol, and Recovery scans.

#### Ketamine

Ketamine is a multi-faceted drug, in terms of both neurophysiology and how it affects subjective experience. Depending on dosage, it can act as a “dissociative” anaesthetic (high dose)^22, 24, 75, 76^ or as an “atypical psychedelic” (at sub-anaesthetic dose)^40–44^. At small doses, it has also found recent use as a fast-acting antidepressant^197, 198^. Both the anaesthetic and psychedelic effects of ketamine are in some respect unusual; unlike widely used anaesthetics like propofol and sevoflurane, ketamine does not exert its anaeshtetic function through agonism of GABA receptors, nor does it recruit sleep-promoting hypothalamic nuclei, which it appears to suppress instead^24, 43^. Likewise, although ketamine does induce psychedelic-like symptoms such as perceptual distortions, vivid imagery and hallucinations, like classic psychedelics, it also induces prominent dissociative symptoms of disembodiment ^43, 64, 199–201^. Its psychedelic action is not mediated by the serotonin 2A receptor, on which classic psychedelics operate^46, 49^: although its precise mechanisms of action are yet to be fully elucidated, ketamine appears to be primarily an antagonist of NMDA and HCN1 receptors; however, evidence suggests that cholinergic, aminergic, and opioid systems may also play modulatory _roles44,45,50._

##### Anaesthetic ketamine dataset: Recruitment

The anaesthetic ketamine data used here have been published before ^75^, and we refer the reader to the original publication for details. The original study was approved by approval by the ethics committee of the Medical school of the university of Liege (University Hospital, Liege, Belgium), registered at eudract 2010-023016-13. 14 right-handed volunteers were recruited via advertisements in an Internet forum (5 women; median age [range], 25 [19 to 31] years). Each participant provided written informed consent to participation, and underwent medical interview and physical examination before their participation.

##### Anaesthetic ketamine dataset: Study protocol

The volunteers were requested to fast for at least 6 h from solids and 2 h from liquids before the experimental session. After structural MR image acquisition, subjects were removed from the MRI scanner, and 64 electroencephalogram (EEG) scalp electrodes were placed to allow for simultaneous EEG-fMRI. Here, we only focus on the functional MRI data.

An 18-gauge intravenous catheter (BD Insyte-W; Becton Dickinson Infusion Therapy Systems inc., USA) was then placed into a vein of the left forearm and infused using normal saline at a rate of 20 ml/h. The intravenous line served for ketamine infusion and eventual administration of rescue medications. a 20-gauge arterial catheter (Arrow International Inc., USA) was also placed into the left radial artery, under strict sterile conditions and after performing local anesthesia with 3 ml of 1% lidocaine. This catheter was equipped with a monitoring set (TruWave, Edwards Lifesciences, Dominican Republic) and served for arterial blood sampling and gas analysis. standard MRI compatible anaesthesia monitoring (Magnitude 3150M; Invivo Research, inc., USA) was also placed to allow continuous monitoring and recording of the electrocardiogram, heart rate, blood pressure, pulse oxymetry (spO2), and breathing frequency throughout the scanning and recovery periods. Through a loosely fitting plastic facemask, additional oxygen at a rate of 5 l/min was provided to volunteers, whose breathing always remained spontaneous. One certified anaesthesiologist and one neurologist were present throughout the experiment. After setting all needed equipment and monitoring, the volunteers were comfortably installed in the MRI tray. The most comfortable supine position attainable was sought to avoid painful stimulation related to position. all volunteers wore earplugs to attenuate noise and earphones to allow communication with investigators; one investigator remained in the MRI scan room at all times.

Ketamine was administered using a computer-controlled intravenous infusion device composed of a separate laptop computer. A 50-ml syringe was filled with normal saline containing racemic ketamine (Ketalar, Pfizer Ltd., Turkey) at a concentration of 10mg/ml. The pharmacokinetic model used to drive the pump was the domino model, which has been demonstrated to have acceptable predictive performance. This system commands infusion pump rates to allow targeting precise effect-site and plasma concentrations of ketamine, based on several biometric parameters. For each change in ketamine concentration, a 5-min equilibration period was allowed after reaching the target, to permit equilibration of ketamine concentration between body compartments. The depth of sedation was assessed using the Ramsay Scale and the University of Michigan Sedation Scale. Each evaluation took place immediately before and after each fMRi data acquisition sequence. Volunteers were asked to strongly squeeze the hand of the investigator, and the command was repeated twice. For that purpose, and for close watch of the volunteer, an investigator continuously stayed inside the MRI room.

A first fMRI data acquisition was performed in the absence of any infusion of ketamine. Ketamine infusion was then started, and its target concentration was increased by steps of 0.5 μg/ml until a level of sedation corresponding to RS 3 to 4 or UMSS 1 to 2 was reached (light sedation, S1). After the 5-min equilibration period, a novel sequence of data acquisition occurred, consisting of the same sequence of events as during W1. Ketamine target concentration was then further increased by steps of 0.5 μg/ml until RS 5 to 6 or UMSS 4 (deep sedation [s2]), and the same sequence of data acquisition was again performed. Because ketamine has a long elimination half-life and to limit time spent in the fMRI scanner for the volunteer, the temporal order of those clinical states was not randomized. For the same reason, a recovery experimental condition could not be achieved. After those acquisitions, the infusion of ketamine was stopped, and the subject was removed from the fMRI scanner to allow for comfortable recovery. The presence of dreaming during ketamine infusion was checked through a phone call at distance from the experimental session. Here, we only consider the awake and deep sedation scans.

##### Anaesthetic ketamine dataset: MRI Data Acquisition

MRI data were acquired on a 3T Siemens Allegra scanner (Siemens AG, Germany; Echo Planar Imaging sequence using 32 slices; repetition time = 2,460 ms; echo time = 40 ms; field of view = 220 mm; voxel size = 3.45 × 3.45× 3 mm; and matrix size = 64 × 64× 32). A high-resolution structural T1 image was acquired in each volunteer at the beginning of the whole experiment for coregistration to the functional data.

A total of 6 participants had to be excluded from the study and further data analysis because of excessive agitation and movements (5 subjects) or voluntary withdrawal (1 subject), leaving 8 for analysis^75^.

##### Psychedelic ketamine dataset: Recruitment

The sub-anaesthetic (“psychedelic”) ketamine data included in this study have been published before^64^, and we refer the reader to the original publication for details. Briefly, a total of 21 participants (10 males; mean age 28.7 years, SD = 3.2 years) were recruited via advertisements placed throughout central Cambridge, UK^64^. All participants underwent a screening interview in which they were asked whether they had previously been diagnosed or treated for any mental health problems and whether they had ever taken any psychotropic medications. Participants reporting a personal history of any mental health problems or a history of any treatment were excluded from the study. All participants were right-handed, were free of current of previous psychiatric or neurological disorder or substance abuse problems, and had no history of cardiovascular illness or family history of psychiatric disorder/substance abuse. The study was approved by the Cambridge Local Research and Ethics Committee, and all participants provided written informed consent in accordance with ethics committee guidelines.

##### Psychedelic ketamine dataset: Study protocol

Participants were scanned (resting-state functional MRI and anatomical T1) on two occasions, separated by at least 1 week. On one occasion, they received a continuous computer-controlled intravenous infusion of a racemic ketamine solution (2 mg/ml) until a targeted plasma concentration of 100 ng/ml was reached. This concentration was sustained throughout the protocol. A saline infusion was administered on the other occasion. Infusion order was randomly counterbalanced across participants. The infusion was performed and monitored by a trained anaesthetist (R.A.) who was unblinded for safety reasons, but who otherwise had minimal contact with participants. At all other times, participants were supervised by investigators blinded to the infusion protocol. The participants remained blinded until both assessments were completed. Bilateral intravenous catheters were inserted into volunteers’ forearms, one for infusion, and the other for serial blood sampling. A validated and previously implemented^202^ three-compartment pharmacokinetic model was used to achieve a constant plasma concentration of 100 ng/ml using a computerized pump (Graseby 3500, Graseby Medical, UK). The infusion continued for 15 min to allow stabilization of plasma levels. Blood samples were drawn before and after the resting fMRI scan and then placed on ice. Plasma was obtained by centrifugation and stored at −70 °C. Plasma ketamine concentrations were measured by gas chromatography–mass spectrometry.

##### Psychedelic ketamine dataset: MRI Data Acquisition

All MRI and assessment procedures were identical across assessment occasions. Scanning was performed using a 3.0 T MRI scanner (Siemens Magnetom, Trio Tim, Erlangen, Germany) equipped with a 12-channel array coil located at the Wolfson Brain Imaging Centre, Addenbrooke’s Hospital, Cambridge, UK. T2*-weighted echo-planar images were acquired under eyes-closed resting-state conditions. Participants were instructed to close their eyes and let the minds wander without going to sleep. Subsequent participant debriefing ensured that no participants fell asleep during the scan. Imaging parameters were: 3x3x3.75mm voxel size, with a time-to-repetition (TR) of 2000 ms, time-to-echo (TE) of 30 ms, flip angle of 781 in 64x64 matrix size, and 240mm field of view (FOV). A total of 300 volumes comprising 32 slices each were obtained. In addition, high- resolution anatomical T1 images were acquired using a three-dimensional magnetic-prepared rapid gradient echo (MPPRAGE) sequence. In all, 176 contiguous sagittal slices of 1.0mm thickness using a TR of 2300 ms, TE of 2.98 ms, flip angle of 91, and a FOV of 256mm in 240x256 matrix were acquired with a voxel size of 1.0mm^3^. One participant was excluded due to excessive movement, resulting in a final sample of N=20 subjects.

#### LSD

LSD (lysergic acid diethylamide) is perhaps the best-known among classic psychedelics, inducing a powerful state of altered consciousness with subjective experiences including hallucinations and “ego dissolution”^46, 48, 49^. Substantial work in humans and animals has demonstrated that LSD influences neuromodulation, having affinity for multiple receptors, primarily serotonergic (5-HT2A, 5-HT1A/B, 5- HT6, 5-HT7) and dopaminergic (D1 and D2 receptors)^46, 48, 52, 203, 204^.

The main neural and subjective effects of LSD originate from its agonism of the 5- HT2A receptor: both effects are abolished by pre-treatment with the non-selective 5HT2 antagonist ketanserin, which has highest affinity for the 5HT2A receptor^171, 205^. In humans, functional connectivity under LSD shows significant correspondence with the spatial distribution of the 5HT2A receptor^206^. Providing evidence for a mechanistic role, both PET maps and transcriptomic maps of the 5HT2A receptor (but not other serotonin receptors) have been shown to improve the ability of computational models to recapitulate the effects of LSD on brain activity and connectivity, as measured by fMRI^72, 73, 178^. Therefore, pharmacological and in silico evidence converge towards the central role of the 5HT2A receptor for LSD’s ability to alter consciousness and its neural underpinnings – although other receptors have also been shown to play an auxiliary role^52^.

##### LSD dataset: Recruitment

The LSD data employed here have been extensively published, and we refer to the original publication for details^60^. Briefly, collection of these data^60^ was approved by the National Research Ethics Service Committee London–West London and was conducted in accordance with the revised declaration of Helsinki (2000), the International Committee on Harmonization Good Clinical Practice guidelines and National Health Service Research Governance Framework. Imperial College London sponsored the research, which was conducted under a Home Office license for research with schedule 1 drugs. All participants were recruited via word of mouth and provided written informed consent to participate after study briefing and screening for physical and mental health. The screening for physical health included electrocardiogram (ECG), routine blood tests, and urine test for recent drug use and pregnancy. A psychiatric interview was conducted and participants provided full disclosure of their drug use history. Key exclusion criteria included: < 21 years of age, personal history of diagnosed psychiatric illness, immediate family history of a psychotic disorder, an absence of previous experience with a classic psychedelic drug (e.g. LSD, mescaline, psilocybin/magic mushrooms or DMT/ayahuasca), any psychedelic drug use within 6 weeks of the first scanning day, pregnancy, problematic alcohol use (i.e. > 40 units consumed per week), or a medically significant condition rendering the volunteer unsuitable for the study. Twenty healthy volunteers with previous experience using psychedelic drugs were scanned.

##### LSD dataset: Study protocol

Volunteers underwent two scans, 14 days apart. On one day they were given a placebo (10-mL saline) and the other they were given an active dose of LSD (75 μg of LSD in 10-mL saline). The order of the conditions was balanced across participants, and participants were blind to this order but the researchers were not. Participants carried out VAS-style ratings via button-press and a digital display screen presented after each scan, and the 11-factor altered states of consciousness (ASC) questionnaire was completed at the end of each dosing day^60^. All participants reported marked alterations of consciousness under LSD.

The data acquisition protocols were described in detail in the original publication ^60^, so we will only describe them in brief here. The infusion (drug/placebo) was administered over 2 min and occurred 115 min before the resting-state scans were initiated. After infusion, subjects had a brief acclimation period in a mock MRI scanner to prepare them for the experience of being in the real machine. ASL and BOLD scanning consisted of three seven-minute eyes closed resting state scans. The ASL data were not analysed for this study, and will not be discussed further.

##### LSD dataset: MRI Data Acquisition

The first and third scans were eyes-closed, resting state without stimulation, while the second scan involved listening to music; however, this scan was not used in this analysis. The precise length of each of the two BOLD scans included here was 7:20 minutes. For the present analysis, these two scans were concatenated together in time. Imaging was performed on a 3T GE HDx system. High-resolution anatomical images were acquired with 3D fast spoiled gradient echo scans in an axial orientation, with field of view = 256x256x192 and matrix = 256x256x129 to yield 1mm isotropic voxel resolution. TR/TE = 7.9/3.0ms; inversion time = 450ms; flip angle = 20. BOLD-weighted fMRI data were acquired using a gradient echo planer imaging sequence, TR/TE = 2000/35ms, FoV = 220mm, 64x64 acquisition matrix, parallel acceleration factor = 2, 90 flip angle. Thirty five oblique axial slices were acquired in an interleaved fashion, each 3.4mm thick with zero slice gap (3.4mm isotropic voxels). One subject aborted the experiment due to anxiety and four others were excluded for excessive motion (measured in terms of frame-wise displacement), leaving 15 subjects for analysis (11 males, 4 females; mean age 30.5 years, SD = 8.0 years)^60^.

#### Psilocybin

Psilocybin (4-phosphoryloxy-N,N-dimethyltryptamine), a prodrug of psilocin (4-OH- N,N-dimethyltrayptamine), is a classic psychedelic, the active compound of “magic mushrooms” of the *Psilocybe* family. Although its psychedelic effects are exerted via agonism of the serotonin 2A receptor^16, 207–210^, psilocin also has demonstrated affinity for additional receptors, in particular serotonin 1A and 2C receptors^211^.

##### Psilocybin dataset: Recruitment

Data acquisition for this dataset is described in detail previously^61^, and will only be summarised here. Volunteers were at least 21 years of age, with no personal or family history of a major psychiatric disorder, no substance dependence, no cardiovascular disease, and no history of adverse response to a psychedelic drug yielded datasets for nine participants. All subjects had used psilocybin at least once before, but not within 6 weeks of the study. The study was approved by a National Health Service research ethics committee and all participants gave informed consent to participate in the study.

##### Psilocybin dataset: Study protocol

In brief, fifteen health volunteers underwent two MRI scanning sessions at least 14 days apart. In each session, subjects were injected. with either psilocybin (2 mg dissolved in 10 mL of saline, 60-s intravenous injection) or a placebo (10 mL of saline, 60-s i.v. injection) in a counterbalanced design. The infusions began exactly 6 min after the start of the 12-min fMRI scans and lasted 60s. The subjective effects of psilocybin were felt almost immediately after injection and sustained for the remainder of the scanning session. At the end of each session, subjects were asked to rate the overall intensity of their subjective experience under the drug (or placebo) and to comment on their wakefulness level throughout the scan. As expected, all participants rated the subjective effects of psilocybin (mean intensity = 6.9/10 ± 2.6) as much stronger than placebo (mean intensity = 0.4/10 ± 0.6) and none of the subjects reported falling asleep during either scanning session. The 5 minutes of post-infusion data were used for the present analysis.

##### Psilocybin dataset: MRI Data Acquisition

Neuroimaging data were acquired using a 3T GE HDx MRI system. Anatomical scans were performed before each functional scan and thus prior to administering either the drug or placebo. Structural scans were collected using a 3D fast spoiled gradient echo scans in an axial orientation, with field of view = 256 × 256 × 192 and matrix = 256 × 256 × 192 to yield 1 mm isotropic voxel resolution (repetition time/echo time TR/TE = 7.9/3.0 ms; inversion time = 450 ms; flip angle = 20). BOLD-weighted fMRI data were acquired at 3T using a gradient echo EPI sequence, TR/TE 3000/35 ms, field-of-view = 192 mm, 64 × 64 acquisition matrix, parallel acceleration factor = 2, 90° flip angle. Fifty-three oblique axial slices were acquired in an interleaved fashion, each 3 mm thick with zero slice gap (3 × 3 × 3-mm voxels). Following the same exclusion criteria for motion described above for the LSD dataset, N=9 subjects were kept for analysis (seven men; age, 32 ± 8.9 SD years of age).

#### DMT

The tryptamine N,N-di-methyltryptamine (DMT) is a fast-acting mind-altering drug, capable of inducing an immersive state of altered consciousness, with vivid and detailed visual hallucinations. It is found endogenously in trace amounts in the human body^46, 212, 213^, but it is orally inert and therefore primarily studied as intravenous injection, whereupon its effects have very rapid onset (2-5 minutes) and offset, effectively fading within 30 minutes of administration^214–217^. Pharmacologically, DMT binds to sigma-1 and serotonin (particularly 2A and 2C) receptors, but also dopamine D1 and alpha-adrenergic receptors^218–221^.

##### DMT dataset: Recruitment

The original DMT study (Timmermann et al., under review) was approved by the National Research Ethics (NRES) Committee London – Brent and the Health Research Authority and was conducted under the guidelines of the revised Declaration of Helsinski (2000), the International Committee on Harmonisation Good Clinical Practices guidelines, and the National Health Service Research Governance Framework. Imperial College London sponsored the research, which was conducted under a Home Office license for research with Schedule 1 drugs. An initial visit was focused on assessing physical and mental health to ensure suitability, and participants provided written informed consent. In total, 20 participants completed all study visits (7 female, mean age = 33.5 years, SD = 7.9).

##### DMT dataset: Study protocol

This was a single-blind, placebo-controlled, counter-balanced design. Volunteers participated in two testing days, 2 weeks apart. On each testing day, volunteers (after testing for drugs of abuse) were involved in 2 separate scanning sessions. In this initial session (task-free) they received intravenous (IV) administration of either placebo (saline) or DMT (in fumarate form) in a counter-balanced order (half of the participants received placebo and the other half received DMT). This first session always consisted of continuous resting-state scans which lasted 28 minutes with DMT/placebo administered at the end of the 8th minute and scanning was over 20 minutes after injection. Participants lay in the scanner with their eyes closed (an eye mask was used to prevent eyes-opening). Following the scanning procedure, participants were interviewed and completed questionnaires designed to assess the subjective effects experienced during the scan. Here, we focused on the 8 minutes post-infusion, corresponding to peak DMT experience.

##### DMT dataset: MRI Data Acquisition

MR images were acquired in a 3T MR scanner (Siemens Magnetom Verio syngo MR B17) using a 12-channel head coil for compatibility with EEG acquisition. Functional imaging was performed using a T2*-weighted BOLD sensitive gradient echo planar imaging sequence (repetition time (TR) = 2000ms, echo time (TE) = 30ms, acquisition time (TA) = 28.06 mins, flip angle (FA) = 80°, voxel size = 3.0 x 3.0 x 3.0mm3, 35 slices, interslice distance = 0mm. Whole-brain T1-weighted structural images were also acquired. EEG was also acquired, but here we only focus on the fMRI data. Seven out of 20 participants were discarded from analyses due to excessive movement, or only completing one session, leaving 13 subjects for analysis.

#### Ayahuasca

The Amazonian beverage ayahuasca is typically used in shamanic religious rituals, where it is obtained as a tea made from two plants: *Psychotria viridis* and *Banisteriopsis caapi*. *Psychotria viridis* contains DMT, which binds to sigma-1 and serotonin (particularly 2A) receptors^218, 219^. *Banisteriopsis caapi* contains beta-carboline alkaloids, notably harmine, tetrahydroharmine (THH), and harmaline. As potent monoamine oxidase inhibitors (MAOI), harmine and harmaline prevent the degradation of DMT by liver MAO that would otherwise render it orally inert; additionally, they also increase levels of monoamine neurotransmitters; and THH acts as a mild selective serotonin reuptake inhibitor and a weak MAOI^222–224^. As for other classic psychedelics, engagement of the 5-HT2A receptor appears to be a necessary condition for the brain and subjective effects of ayahuasca to manifest ^225^.

##### Ayahuasca dataset: Recruitment

The ayahuasca data that we used have been published before^62, 144^, and we refer the reader to the original publication for details. Briefly, data were obtained from 9 healthy right-handed adult volunteers (mean age 31.3, from 24 to 47 years), all who were experienced users of Ayahuasca with at least 5 years use (twice a month) and at least 8 years of formal education. The experimental procedure was approved by the Ethics and Research Committee of the University of São Paulo at Ribeirão Preto (process number 14672/2006). Written informed consent was obtained from all volunteers, who belonged to the Santo Daime religious organisation. All experimental procedures were performed in accordance with the relevant guidelines and regulations. Volunteers were not under medication for at least 3 months prior to the scanning session and were abstinent from caffeine, nicotine and alcohol prior to the acquisition. They had no history of neurological or psychiatric disorders, as assessed by DSM-IV structured interview71. Subjects ingested 120–200 mL (2.2 mL/kg of body weight) of Ayahuasca known to contain 0.8 mg/mL of DMT and 0.21 mg/mL of harmine. Harmaline was not detected via the chromatography analysis, at the threshold of 0.02 mg/mL7.

##### Ayahuasca dataset: Study protocol

Volunteers underwent two distinct fMRI scanning sessions: (i) before and (ii) 40 minutes after Ayahuasca intake, when the subjective effects become noticeable (the volunteers drank 2.2 mL/kg of body weight and the Ayahuasca contained 0.8 mg/mL of DMT and 0.21 mg/mL of harmine). In both cases, participants were instructed to close their eyes and remain awake and at rest, without performing any task.

##### Ayahuasca dataset: MRI Data Acquisition

The fMRI images were obtained in a 1.5 T scanner (Siemens, Magneton Vision), using an EPI-BOLD like sequence comprising 150 volumes, with the following parameters: TR = 1700 ms; TE = 66 ms; FOV = 220 mm; matrix 64 ×64; voxel dimensions of 1.72 mm × 1.72 mm × 1.72 mm. Whole brain high resolution T1- weighted images were also acquired (156 contiguous sagittal slices) using a multiplanar reconstructed gradient-echo sequence, with the following parameters: TR = 9.7 ms; TE = 44 ms; flip angle 12°; matrix 256 × 256; FOV = 256 mm, voxel size = 1 mm× 1 mm× 1 mm. Data from one volunteer were excluded from analyses due to acquisition limitations resulting in incomplete brain coverage. The final dataset included 8 subjects.

#### MDMA

3,4-methylenedioxymethamphetamine (MDMA) combines the subjective effects of a stimulant and a psychedelic, inducing powerful euphoria and prosociality, but also mild visual hallucinations^226–228^. It inhibits reuptake of noradrenaline, dopamine, and serotonin by acting on their respective transporters, and it also stimulates their release – with preferential effects on 5-HT mediating at least in part its induction of positive mood^226–228^.

##### MDMA dataset: Recruitment

The MDMA dataset has been published before^63, 229^, and we refer the reader to the original publications for details. The study was approved by the National Research Ethics Service West London Research Ethics Committee, Joint Compliance and Research Office of Imperial College London, Research Ethics Committee of Imperial College London, Head of the Department of Medicine of Imperial College London, Imanova Centre for Imaging Science, and Faculty of Medicine of Imperial College London. The study was conducted in accordance with Good Clinical Practice guidelines. A Home Office Licence was obtained for the storage and handling of a Schedule 1 drug. Imperial College London sponsored the research.

The study included 25 healthy participants (mean age, 34±11 years; 7 females) with at least one previous experience with MDMA. None of the participants had used MDMA for at least 7 days or other drugs for at least 48 hours, which was confirmed by a urine screen. An alcohol breathalyzer test confirmed that none of the participants had recently consumed alcohol. Volunteers were screened for good physical and mental health, and magnetic resonance imaging compatibility. All subjects were deemed physically and mentally healthy, and none had any history of drug or alcohol dependence or diagnosed psychiatric disorder.

##### MDMA dataset: Study protocol

The study design was within-subjects, and placebo-controlled. Participants were scanned twice, once after MDMA (100 mg encapsulated MDMA-HCl) and once after placebo (encapsulated ascorbic acid/vitamin-C), in a double-blind, randomized, counterbalanced order. The MDMA and placebo study days were separated by 7 days, and on each occasion, participants underwent two arterial spin labelling (ASL; not included in the present analysis) and two resting-state BOLD fMRI scans within a 90-minute scan session. Biochip Array Technology (Randox Laboratories Ltd., Co., Antrim, United Kingdom) was used to detect MDMA from plasma samples obtained shortly after each participant’s MDMA scanning session (i.e., 2 hours after capsule ingestion).

The first resting-state BOLD scan took place 60 minutes after capsule ingestion and the second resting-state BOLD scan occurred 113 minutes after capsule ingestion. Participants relaxed with their eyes closed during the ASL and BOLD resting state scans. Peak subjective effects were reported ∼100 minutes post administration of MDMA, consistent with the plasma t-max of MDMA. Therefore, here we focus on the second MDMA and placebo rs-fMRI scans.

##### MDMA dataset: MRI Data Acquisition

MR images were acquired on a 3T Siemens Tim Trio (Siemens Healthcare, Erlangen, Germany) using a 32-channel phased array head coil. Anatomical reference images were acquired using the ADNI-GO recommended MPRAGE parameters (1 mm isotropic voxels, TR = 2300 ms, TE = 2.98 ms, 160 sagittal slices, 256 x 256 in-plane resolution, flip angle = 9 degrees, bandwidth = 240 Hz/pixel, GRAPPA acceleration = 2). T2*-weighted echo-planar images (EPI) were acquired for the resting state BOLD functional scan using 3 mm isotropic voxels in a 192 mm in-plane FOV, TR = 2 s, echo time = 31 ms, 80 degree flip angle, 36 axial slices in each TR, bandwidth = 2298 Hz/pixel, and a GRAPPA acceleration of 2. Each rs-fMRI scan lasted for 6 minutes, corresponding to 180 functional volumes.

#### Modafinil

Modafinil is a wakefulness promoting drug used for the treatment of sleep disorders such as narcolepsy (under the commercial name Provigil), as well as finding use as a cognitive enhancer for attention and memory^6, 7, 65^, and to combat the cognitive symptoms of Attention Deficit/Hyperactivity Disorder (ADHD), and mood disorders, owing to its lower addiction risk in comparison with amphetamine-like psychostimulants^6–9, 230^. This drug has a broad neurotransmitter profile: it acts as a blocker of the dopamine and noradrenaline transporters, as well as modulating locus coeruleus noradrenergic firing, and acting on the wake-promoting hypothalamic neuropeptide orexin to activate the histamine system; it also influences both the glutamate and GABA systems^8, 231–235^.

##### Modafinil dataset: Recruitment

The modafinil dataset has been published before^65, 236^. As reported in the original publication, the study was approved by the ethics committee of University of Chieti (PROT 2008/09 COET on 14/10/2009) and conducted in accordance with the Helsinki Declaration. The study design was explained in detail and written informed consent was obtained from all participants involved in our study. Recruitment was performed throughout February 2011, drug/placebo administration and fMRI acquisitions started on March 2011, went on until January 2012, and the study was completed with the last fMRI session in January 2012. After securing financial coverage for costs related to the analysis of the study, the trial was registered on 10/09/2012 (NCT01684306http://clinicaltrials.gov/ct2/show/NCT01684306). After obtaining registration, the double-blind study was opened and analyzed.

This dataset was obtained from the OpenfMRI database. Its accession number is ds000133. A total of twenty six young male right-handed (as assessed by the Edinburgh Handedness inventory) adults (age range: 25–35 years) with comparable levels of education (13 years) were enrolled. All subjects had no past or current signs of psychiatric, neurological or medical (hypertension, cardiac disorders, epilepsy) conditions as determined by the Millon test and by clinical examination. Subjects showing visual or motor impairments were excluded as well as individuals taking psychoactive drugs or having a history of alcohol abuse. All volunteers were instructed to maintain their usual amount of nicotine and caffeine intake and avoid alcohol consumption in the 12h before the initiation of the study.

##### Modafinil dataset: Study protocol

Study subjects received, in a double blind fashion, either a single dose of modafinil (100 mg) (modafinil group; N=13) or a placebo (placebo group; N=13) pill identical to the drug. Randomization of study subjects was obtained by means of a random number generator. Here, we only considered data from the modafinil group. The day after drug/placebo assumption, subjects were asked about perceived side effects and, in particular, sleep disturbances. All but one reported no modafinil-induced side- effects or alterations in the sleep-wake cycle. Rs-fMRI BOLD data were separated in three runs lasting four minutes each followed by high resolution T1 anatomical images. Two scanning sessions took place: one before ingesting the drug/placebo, and one 3 hours later, to account for pharmacokinetics. Subjects were asked to relax while fixating the central point in the middle of a grey-background screen that was projected on an LCD screen and viewed through a mirror placed above the subject’s head. Subject head was positioned within an eight-channel coil and foam padding was employed to minimize involuntary head movements.

##### Modafinil dataset: MRI Data Acquisition

BOLD functional imaging was performed with a Philips Achieva 3T Scanner (Philips Medical Systems, Best, The Netherlands), using T2*-weighted echo planar imaging (EPI) free induction decay (FID) sequences and applying the following parameters: TE 35 ms, matrix size 64x64, FOV 256 mm, in-plane voxel size 464 mm, flip angle 75 degrees, slice thickness 4 mm and no gaps. 140 functional volumes consisting of 30 transaxial slices were acquired per run with a volume TR of 1,671 ms. High- resolution structural images were acquired at the end of the three rs-fMRI runs through a 3D MPRAGE sequence employing the following parameters: sagittal, matrix 256x256, FOV 256 mm,slice thickness 1 mm, no gaps, in-plane voxel size 1 mm x 1 mm,flip angle 12 degrees, TR = 9.7 ms and TE = 4 ms. Two subjects from the modafinil group were excluded from analysis due to acquisition limitations, leaving N=11 subjects for analysis.

#### Methylphenidate

Methylphenidate is used as a cognitive enhancer to treat the cognitive symptoms of ADHD and narcolepsy (under the name Ritalin)^4, 11^. Pharmacologically, it inhibits the reuptake of both dopamine and noradrenaline by blocking their transporters; although yet to be conclusively confirmed, there is also in vitro evidence suggesting an additional minor affinity of methylphenidate for the 5-HT1A receptor^237–243^.

##### Methylphenidate dataset: Recruitment

The methylphenidate dataset used here has been published before^10, 51^. Unlike the other datasets included in this study, the methylphenidate data were not obtained from healthy controls, but rather from a cohort of patients suffering from traumatic brain injury (TBI). Volunteers with a history of moderate to severe traumatic brain injury (inclusion criteria: age 18–60 years and not recruited to more than three research studies within the calendar year) were referred from the Addenbrooke’s Neurosciences Critical Care Unit Follow-Up Clinic, Addenbrooke’s Traumatic Brain Injury Clinic and The Royal London Hospital Intensive Care Unit (see the original publication for details of patient injuries). The patients were sent a written invitation to take part in the study. All volunteers gave written informed consent before participating in the study.

Thirty-eight volunteers were recruited to the study; 17 (12 male, 5 female) into the TBI arm of the study and 21 (13 male, 8 female) into the healthy control (HC) arm of the study. Exclusion criteria included National Adult Reading Test (NART) <70, Mini Mental State Examination (MMSE) <23, left-handedness, history of drug/alcohol abuse, history of psychiatric or neurological disorders, contraindications for MRI scanning, medication that may affect cognitive performance or prescribed for depression, and any physical handicap that could prevent the completion of testing.

Our sample contains mostly patients with diffuse axonal injuries and small lesions. The patients were at least 6 months post TBI. Four sustained moderate TBI with a score of between 9 and 12 on the Glasgow Coma Scale (GCS) and 11 sustained severe TBI with a GCS score of 8 or below on presentation. The mean age of the patient group was 36 years (±13 years).

##### Methylphenidate dataset: Study protocol

The study consisted of two visits (separated by 2–4 weeks) for both groups of participants. The TBI volunteers were randomly allocated in a Latin square design to receive one of the two interventions on the first visit (a placebo tablet or 30 mg tablet of methylphenidate), and the alternate intervention on the second visit. The decision to use 30 mg of methylphenidate was based on comparable doses used in previous studies in healthy participants, as well as NICE guidelines for medication in adults (www.nice.org.uk) which stipulate that when methylphenidate is titrated for side effects and responsiveness in each individual subject, the dose should range from a minimum of 15 mg to a maximum dose of 100 mg. As the dose of methylphenidate was not calculated by the participant’s body weight, an interventional dose at the lower end of the dose range was chosen. After a delay of 75 min to ensure that peak plasma levels of methylphenidate were reached, the volunteers completed an MRI scan which included both fMRI and structural image acquisition. The healthy controls attended their two fMRI assessments at the same time interval as the patients, but without any pharmacological intervention. Therefore, here we only considered the patient data.

##### Methylphenidate dataset: MRI Data Acquisition

MRI data were acquired on a Siemens Trio 3-Tesla MR system(Siemens AG, Munich, Germany). MRI scanning started with the acquisition of a localizer scan and was followed by a 3D high resolution MPRAGE image [TR 2,300 ms, Echo Time (TE) 2.98 ms, Flip Angle 9◦, FOV 256×256 mm2]. Diffusion Tensor Imaging (DTI) data (63non-collinear directions,b=1,000 s/mm2with one volume acquired without diffusion weighting (b=0), echo time 106ms, repetition time 1,700 ms, field of view 192×192 mm,2 mm3isotropic voxels) were also collected to investigate white matter integrity. Here, we did not analyse the DTI data.

Functional imaging data were acquired using an echo-planar imaging (EPI) sequence with parameters TR = 2,000 ms, TE = 30 ms, Flip Angle = 78◦, FOV 192×192 mm2, in-plane resolution 3.0×3.0 mm, 32 slices 3.0 mm thick with a gap of 0.75 mm between slices. Two patients were excluded from the analysis (one patient only attended one of the study sessions and the other had excessive movement artifacts in their fMRI scan), leaving N=15 patients for analysis.

### Functional MRI preprocessing and denoising

#### Preprocessing

To maximise the uniformity between the different datasets included here, we elected to preprocess all datasets using the same pipeline, as opposed to relying on the different pipelines originally employed by the group that collected each dataset. For the same reason, we also applied the same denoising procedure across all datasets.

For each dataset and condition, we applied a standard preprocessing pipeline, which we have previously employed with pharmaco-MRI datasets comprising both anaesthetics and psychedelics^18, 21, 145, 196^, demonstrating its suitability for the present analysis. Preprocessing was performed using the CONN toolbox, version 17f (CONN; http://www.nitrc.org/projects/conn)^244^ based on Statistical Parametric Mapping 12 (http://www.fil.ion.ucl.ac.uk/spm), implemented in MATLAB 2016a. The pipeline involved the following steps: removal of the first 10s, to achieve steady-state magnetization; motion correction; slice-timing correction; identification of outlier volumes for subsequent scrubbing by means of the quality assurance/artifact rejection software *art* (http://www.nitrc.org/projects/artifact_detect); normalisation to Montreal Neurological Institute (MNI-152) standard space (2 mm isotropic resampling resolution), using the segmented grey matter image from each volunteer’s T1-weighted anatomical image, together with an *a priori* grey matter template.

#### Denoising

Denoising was also performed using the CONN toolbox, using the same approach as in our previous publications with pharmaco-MRI datasets^18, 21, 145, 196^, which has also been adopted with similar pharmaco-MRI datasets in publications by independent groups^245^.

Pharmacological agents can induce alterations in physiological parameters (heart rate, breathing rate, motion) or neurovascular coupling^246^. The anatomical CompCor (aCompCor) method removes physiological fluctuations by extracting principal components from regions unlikely to be modulated by neural activity; these components are then included as nuisance regressors^247^. Following this approach, five principal components were extracted from white matter and cerebrospinal fluid signals (using individual tissue masks obtained from the T1-weighted structural MRI images)^245^; and regressed out from the functional data together with six subject-specific realignment parameters (three translations and three rotations) as well as their first-order temporal derivatives; followed by scrubbing of outliers identified by ART, using Ordinary Least Squares regression^244^. Finally, the denoised BOLD signal timeseries were linearly detrended and band-pass filtered to eliminate both low-frequency drift effects and high-frequency noise, thus retaining frequencies between 0.008 and 0.09 Hz.

The step of global signal regression (GSR) has received substantial attention in the literature as a denoising method^248–250^. GSR mathematically mandates that approximately 50% of correlations between regions will be negative^251^; however, the proportion of anticorrelations between brain regions has been shown to vary across states of consciousness, including anaesthesia and psychedelics^18, 145^. Furthermore, recent work has demonstrated that the global signal contains information about states of consciousness, across pharmacological and pathological perturbations^252^. Therefore, in line with ours and others’ previous studies, here we avoided GSR in favour of the aCompCor denoising procedure, which is among those recommended^249^.

### Summarising pharmacological effects on brain function

For each subject at each condition, the denoised regional BOLD signals were parcellated into 100 cortical regions according to the local-global functional parcellation of Schaefer and colleagues^112^. The parcellated regional BOLD signals were then correlated (“functional connectivity”); after removing negative-valued edges, the regional strength of functional connectivity (node strength) was measured for each region. The regional change in FC strength was then quantified for each subject (for the methylphenidate dataset, this was quantified with respect to the mean of controls’ node strength values). Finally, for each dataset, we computed the mean across subjects of the FC strength deltas. Therefore, each pharmacological intervention was summarised as one vector of regional FC strength deltas (Figure 1).

### Receptor maps from Positron Emission Tomography

Receptor densities were estimated using PET tracer studies for a total of 19 receptors and transporters, across 9 neurotransmitter systems, recently made available by Hansen and colleagues at https://github.com/netneurolab/hansen_receptors31. These include dopamine (D1^77^, D2^78–81^, DAT^82^), norepinephrine (NET^83–86)^, serotonin (5-HT1A^87^, 5-HT1B,^87–90, 90–92^ 5-HT2A^93^, 5-HT4^93^, 5-HT6^94, 95^, 5-HTT^93^), acetylcholine (*α*4*β*2^96, 97^, M1^98^, VAChT^99, 100^), glutamate (mGluR5^101, 102^, NMDA^103, 104^), GABA (GABA-A^105^), histamine (H3^106^), cannabinoid (CB1^107–110)^, and opioid (MOR^111^). Volumetric PET images were registered to the MNI-ICBM 152 nonlinear 2009 (version c, asymmetric) template, averaged across participants within each study, then parcellated and receptors/transporters with more than one mean image of the same tracer (5-HT1b, D2, VAChT) were combined using a weighted average. See the dedicated article by Hansen *et al*^31^ for detailed information about each PET dataset and their respective acquisition and limitations.

### Partial Least Squares Analysis

PLS analysis (implemented using MATLAB code from Hansen et al., 2021)^59^ available at https://github.com/netneurolab/hansen_genescognition/tree/master/PLS) was used to relate regional neurotransmitter density to pharmacologically-induced functional connectivity changes. PLS analysis is an unsupervised multivariate statistical technique that decomposes relationships between two datasets (in our case, neurotransmitter density with *n* regions and *r* neurotransmitters, *X_nxr_*, and drug-induced functional connectivity changes, *Y_nxd_* with *n* regions and *d* drugs) into orthogonal sets of latent variables with maximum covariance, which are linear combinations of the original data^119, 120^. In other words, PLS finds components from the predictor variables (100 × 19 matrix of regional neurotransmitter receptor and transporter density scores) that have maximum covariance with the response variables (100 × 15 matrix of regional changes in FC induced by different drugs). The PLS components (i.e., linear combinations of the weighted neurotransmitter density) are ranked by covariance between predictor and response variables, so that the first few PLS components provide a low-dimensional representation of the covariance between the higher dimensional data matrices. Thus, the first PLS component (PLS1) is the linear combination of the weighted neurotransmitter density scores that have a brain expression map that covaries the most with the map of regional FC changes.

This is achieved by z-scoring both data matrices column-wise and applying singular value decomposition on the matrix *Y’X*, such that:

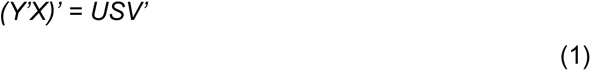

where *U_g×t_* and *V_t×t_* are orthonormal matrices consisting of left and right singular vectors and *S_t×t_* is a diagonal matrix of singular values. The i^th^ columns of *U* and *V* constitute a latent variable, and the i^th^ singular value in *S* represents the covariance between singular vectors. The i^th^ singular value is proportional to the amount of covariance between neurotransmitter density, and drug-induced FC changes captured by the i^th^ latent variable, where the effect size can be estimated as the ratio of the squared singular value to the sum of all squared singular values. In the present study, the left singular vectors (that is, the columns of *U*) represent the degree to which each neurotransmitter contributes to the latent variable and demonstrate the extracted association between neurotransmitter density and drug-induced FC changes (neurotransmitter weights). The right singular vectors (that is, the columns of *V*) represent the degree to which the FC changes contribute to the same latent variable (term weights). Positively weighed neurotransmitters covary with positively weighed drug-induced changes, and negatively weighed neurotransmitters covary with negatively weighed drug-induced changes.

Scores at each brain region for each latent variable can be computed by projecting the original data onto the singular vector weights. Positively scored brain regions are regions that demonstrate the covariance between the prevalence of positively weighted neurotransmitters and positively weighted drug-induced effects (and vice versa for negatively scored brain regions). Loadings for each variable were computed as the Pearson’s correlation between each individual variable’s activity (neurotransmitter density and drug-induced FC changes) and the PLS analysis-derived neurotransmitter score pattern. Squaring the loading (a correlation) equals the percentage variance shared between an original variable and the PLS analysis-derived latent variable. Variables with high absolute loadings are highly correlated to the score pattern, indicating a large amount of shared variance between the individual variable and the latent variable. We confirmed that PLS1 explained the largest amount of variance by testing across a range of PLS components (between 1 and 12) and quantifying the relative variance explained by each component.

### Hierarchical organisation

We quantified the spatial similarity of each pharmacologically-induced pattern of change in FC strength, with several canonical maps of hierarchical brain organisation (“canonical brain hierarchies”) derived from multimodal neuroimaging. We considered the anatomical gradient of intracortical myelination obtained from T1w/T2w MRI ratio^58^; evolutionary cortical expansion obtained by comparing human and macaque^126^; the principal component of variation in gene expression from the Allen Human Brain Atlas transcriptomic database (AHBA; https://human.brain-map.org/), referred to as “AHBA PC1”^54, 59, 127^; the principal component of variation in task activation from NeuroSynth, (https://github.com/neurosynth/neurosynth), an online meta-analytic tool that synthesizes results from more than 15,000 published fMRI studies by searching for high-frequency key words that are published alongside fMRI voxel coordinates, using the volumetric association test maps (referred to as “NeuroSynth PC1”)^54, 59, 128^; the map of cerebral blood flow^54^; the principal gradient of variation in functional connectivity^57^; and a recently derived gradient of regional prevalence of different kinds of information, from redundancy to synergy^129^.

### ENIGMA cortical vulnerability data

Patterns of cortical thickness were collected for the available 11 neurological, neurodevelopmental, and psychiatric disorders from the ENIGMA (Enhancing Neuroimaging Genetics through Meta-Analysis) consortium and the *enigma* toolbox (https://github.com/MICA-MNI/ENIGMA)^130^: 22q11.2 deletion syndrome^131^, attention- deficit/hyperactivity disorder^132^, autism spectrum disorder^133^, idiopathic generalized epilepsy^134^, right temporal lobe epilepsy^134^, left temporal lobe epilepsy^134^, depression^135^, obsessive-compulsive disorder^136^, schizophrenia^137^, bipolar disorder^138^, and Parkinson’s disease^139^. The ENIGMA consortium is a data-sharing initiative that relies on standardized image acquisition and processing pipelines, such that disorder maps are comparable^253^. Altogether, over 21,000 patients were scanned across the thirteen disorders, against almost 26,000 controls. The values for each map are z-scored effect sizes (Cohen’s *d*) of cortical thickness in patient populations versus healthy controls. Imaging and processing protocols can be found at http://enigma.ini.usc.edu/protocols/.

For every brain region, we constructed an 11-element vector of disorder abnormality, where each element represents a disorder’s cortical abnormality at the region. For every pair of brain regions, we correlated the abnormality vectors to quantify how similarly two brain regions are affected across disorders. This results in a region-by-region matrix of “disorder co-susceptibility”^113^.

### Gradients from diffusion map embedding

We used the *BrainSpace* toolbox^140^ (http://brainspace.readthedocs.io) to obtain gradients from diffusion map embedding. A joint network of disease- and drug- co-susceptibility was obtained from the network fusion procedure of Paquola *et al* (2020)^141^, through horizontal concatenation of matrices and production of a node-to-node affinity matrix using row-wise normalised angle similarity.

We then employed diffusion map embedding, a nonlinear manifold learning technique based on the graph Laplacian^140, 254^, to obtain a low dimensional representation of the joint drug- and disorder-susceptibility. A single parameter α controls the influence of the sampling density on the manifold (α = 0, maximal influence; α = 1, no influence). Following extensive previous work using this approach with neuroimaging data^57, 140, 141, 255^, we set α = 0.5, to retain the global relations between data points in the embedded space. The ability to combine global and local geometry differentiates diffusion maps from global-only methods, such as PCA and multidimensional scaling^141^. A small number of components can be identified based on decreasing eigenvalues. The decay of each eigenvector obtained from the diffusion map embedding provides an overall measure of the connectivity between nodes along the axis delineated by its spatial distribution on the brain (gradient).

### Statistical Analysis

The statistical significance of the variance explained by each PLS model was tested by permuting the response variables 1,000 times, while considering the spatial dependency of the data by using spatial autocorrelation-preserving permutation tests, termed spin tests^121–124^. Parcel coordinates were projected onto the spherical surface and then randomly rotated and original parcels were reassigned the value of the closest rotated parcel (10,000 repetitions). The procedure was performed at the parcel resolution rather than the vertex resolution to avoid upsampling the data. In PLS analysis, the spin test is applied to the singular values (or equivalently, the covariance explained) of the latent variables, producing a null distribution of singular values. This is done applying PLS analysis to the original *X* matrix and a spun *Y* matrix. The spin test embodies the null hypothesis that neurotransmitter density and drug-induced FC changes are spatially correlated with each other only because of inherent spatial autocorrelation. The p-value is computed as the proportion of null singular values that are greater in magnitude than the empirical singular values. Thus, these p-values represent the probability that the observed spatial correspondence between neurotransmitter density and drug-induced FC changes could occur by randomly correlating maps with comparable spatial autocorrelation.

Likewise, spatial similarity between brain maps was quantified in terms of Spearman correlation, and statistical significance was assessed against a spin-based null model with preserved spatial autocorrelation, as described above^121–124^.

## Data and code availability

Pharmacological-fMRI data are available upon request from the corresponding authors of the original publications referenced herein. The neurotransmitter receptor and transporter PET maps are available at https://github.com/netneurolab/hansen_receptors. The Allen Human Brain Atlas transcriptomic database is available at https://human.brain-map.org/; NeuroSynth is available at https://github.com/neurosynth/neurosynth. The ENIGMA toolbox and data are available at https://github.com/MICA-MNI/ENIGMA. MATLAB code for Partial Least Squares analysis is freely available at https://github.com/netneurolab/hansen_genescognition/tree/master/PLS.

## Acknowledgements

This work was carried out thanks to support from the Gates Cambridge Trust (OPP 1144) [to AIL]; the Wellcome Trust Research Training Fellowship (grant no. 083660/Z/07/Z), Raymond and Beverly Sackler Studentship, and the Cambridge Commonwealth Trust [to RA]; the Canadian Institute for Advanced Research (CIFAR; grant RCZB/072 RG93193) [to DKM and EAS]; the Cambridge Biomedical Research Centre and NIHR Senior Investigator Awards and the British Oxygen Professorship of the Royal College of Anaesthetists [to DKM]; the Stephen Erskine Fellowship at Queens’ College, Cambridge [to EAS]. BM is supported by the Natural Sciences and Engineering Research Council of Canada (NSERC Discovery Grant RGPIN #017-04265) and Canada Research Chairs Program. JYH is supported by the Helmholtz International BigBrain Analytics & Learning Laboratory, the Natural Sciences and Engineering Research Council of Canada, and Fonds de reserches de Québec. AMO is supported by the Canada Excellence Research Chairs program (215063); LN acknowledges support by the L’Oreal-Unesco for Women in Science Excellence Research Fellowship. LR acknowledges support of the Imperial College President’s Scholarship. PC and NLNA are supported by the Belgian National Funds for Scientific Research (F.R.S-FNRS). PC is also supported by the GIGA-Doctoral School for Health Sciences (University of Liège). NLNA also acknowledges the support of the Human Brain Project. SLS is supported by funds from the Italian Department of Education [Fondo per gli Investimenti della Ricerca di Base (FIRB) 2003; Programmi di Ricerca di Rilevante Interesse nazionale (PRIN) 2008]. RLC-H was supported by the Alex Mosley Charitable Trust and supporters of the Centre for Psychedelic Research during the period of data collection and now holds the Ralph Metzner Distinguished Professorship at UCSF. The original LSD study received support from a Crowd Funding Campaign and the Beckley Foundation, as part of the Beckley-Imperial Research Programme. The MDMA research was supported by funds provided by the British public service broadcast station Channel 4 and was performed as part of the Beckley Foundation–Imperial College research program. The DMT study was funded via a donation by Patrick Vernon. The Beckley Foundation mediated Patrick Vernon’s support. The psilocybin study received support from the Beckley Foundation and financial support from the Neuropsychoanalysis Foundation, Multidisplinary Association for Psychedelic Studies, and the Heffter Research Institute. The Cambridge ketamine study was funded by the Bernard Wolfe Health Neuroscience Fund and the Wellcome Trust. Data acquisition for the sevoflurane dataset was funded by the Departments of Anesthesiology, Neurology, and Neuroradiology of the Klinikum rechts der Isar of the Technical University Munich. The anaesthetic ketamine study was supported by the Belgian Society of Anesthesiology, Resuscitation, Perioperative medicine and Pain management (BeSARPP, Brussels, Belgium), the Belgian National Funds for Scientific Research (Brussels, Belgium), the European Commission (Brussels, Belgium), the James McDonnell Foundation (Saint Louis, Missouri), the European Space Agency (Brussels, Belgium), the Mind Science Foundation (San Antonio, Texas), the French Speaking Community Concerted Research Action (ARC - 06/11 – 340, Brussels, Belgium), the Public Utility Foundation “Université Européenne du Travail” (Brussels, Belgium), “Fondazione Europea di Ricerca Biomedica” (Milan, Italy), and the University and University Hospital of Liege (Liege, Belgium). This work was performed using resources provided by the Cambridge Service for Data Driven Discovery (CSD3) operated by the University of Cambridge Research Computing Service (www.csd3.cam.ac.uk), provided by Dell EMC and Intel using Tier-2 funding from the Engineering and Physical Sciences Research Council (capital grant EP/T022159/1), and DiRAC funding from the Science and Technology Facilities Council (www.dirac.ac.uk). Computing infrastructure at the Wolfson Brain Imaging Centre (WBIC-HPHI) was funded by the MRC research infrastructure award (MR/M009041/1). We acknowledge the contribution of Santo Daime members for volunteering and for providing the Ayahuasca. We also thank members of the Cognition and Consciousness Imaging Group, and of the Network Neuroscience Lab, for many helpful discussions.

## Author Contributions

AIL, JYH, EAS, BM conceived the study. AIL, JYH, EAS, BM designed the methodology and the analysis. AIL analysed the data. JYH, ARDP contributed to data analysis. RA, DKM, EAS, LR, RLCH, CT, DG, AR, RI, DJ, VB, AV, AD, OJ, MAB, NLNA, PC, AEM, DBdA, SLS, AMO, LN were involved in design, execution, and data acquisition and curation for the original studies for which the present data were collected. AIL, EAS, BM wrote the manuscript with feedback from co-authors.

