## Supplementary Information for "Mapping Pharmacologically-induced Functional Reorganisation onto the Brain’s Neurotransmitter Landscape"

### Supplementary Figures

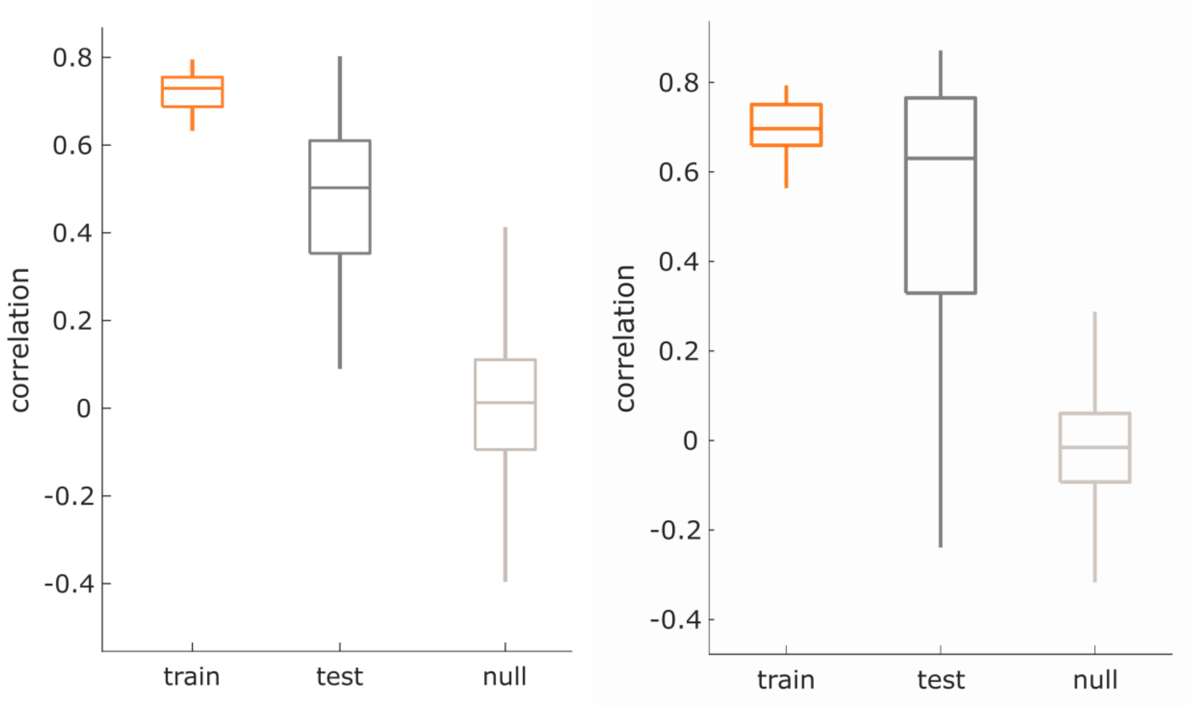


**Figure S1. Distance-dependent cross-validation.** The correlation between drug scores and neurotransmitter scores was cross-validated by constructing the training set with 75% of brain regions closest in Euclidean distance to a randomly chosen source node (red) and with the testing set as the remaining 25% of brain regions (dark grey; 100 repetitions). The out-of-sample mean was significant against a permuted null model (1,000 repetitions; null model shown in light grey; both $p < 0.001$).

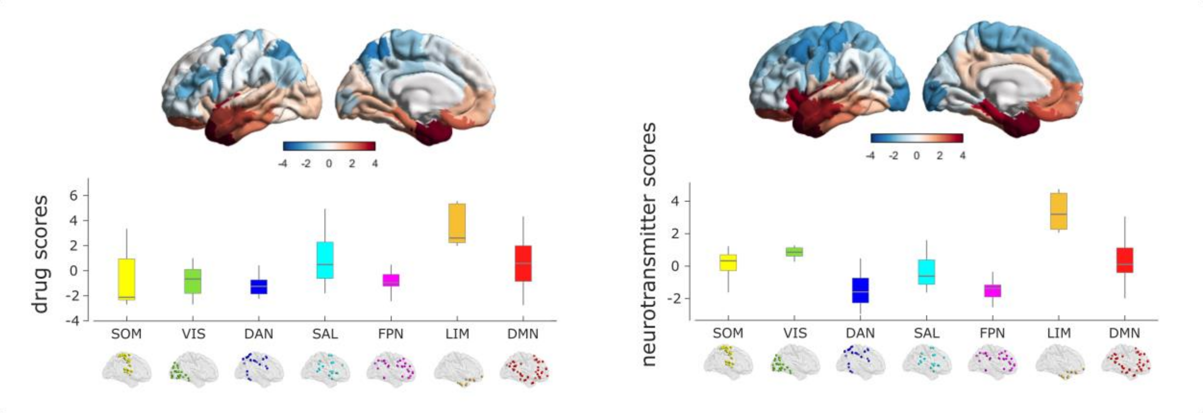


**Figure S2. Association between PLS2 scores and intrinsic resting-state networks from Yeo et al (2011).** Left: drug scores. Right: neurotransmitter scores.

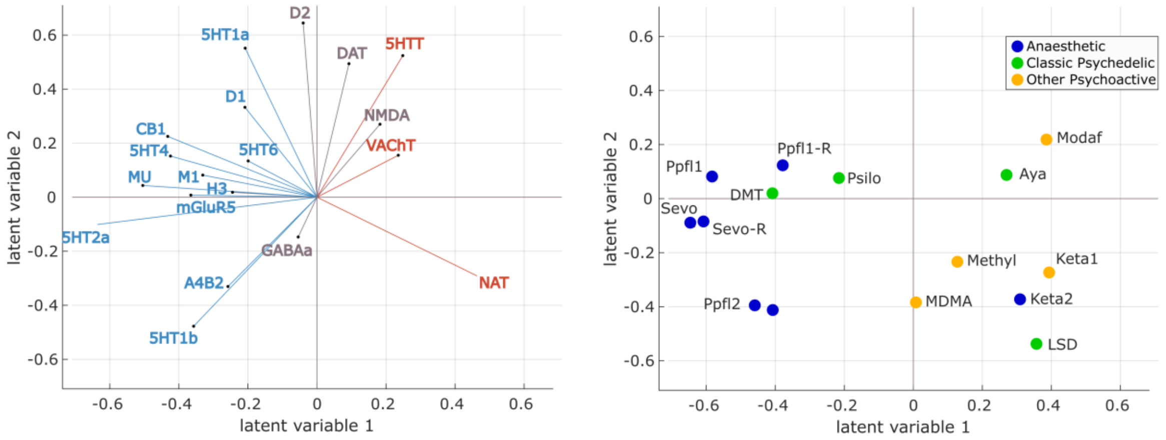


**Figure S3. Neurotransmitters and drugs plotted separately in the same space.** Left: Each neurotransmitter receptor and transporter is represented as a vector in the 2D space of the first two PLS latent variables. Right: Each drug is represented as a point reflecting its projection onto the first two latent variables of the PLS analysis, color-coded based on its effects on subjective experience (anaesthetic, classic psychedelic, or other psychoactive).

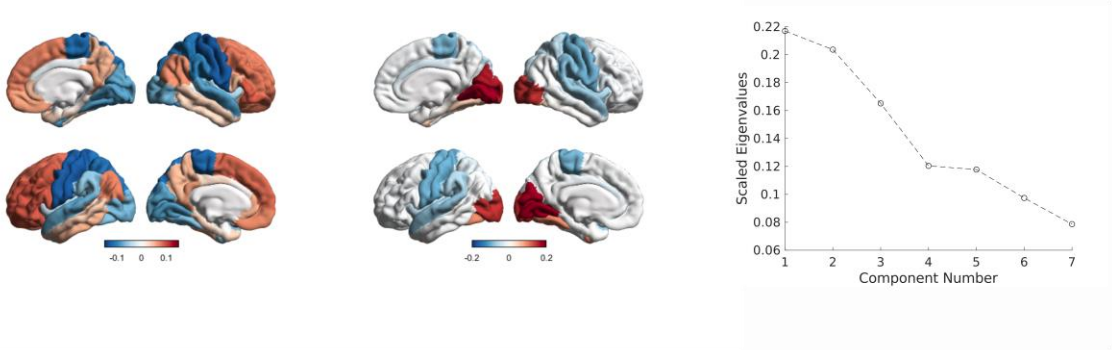


**Figure S4. Gradients of pharmacological susceptibility.** We found that the two principal gradients obtained from diffusion map embedding of the matrix of pharmacological co-susceptibility correspond to the principal components of FC variation: one distinguishing unimodal from transmodal association cortices (left); and the second distinguishing between sensorimotor and visual cortices (middle). Right: eigenvalues as a function of component number, for the nonlinear embedding.

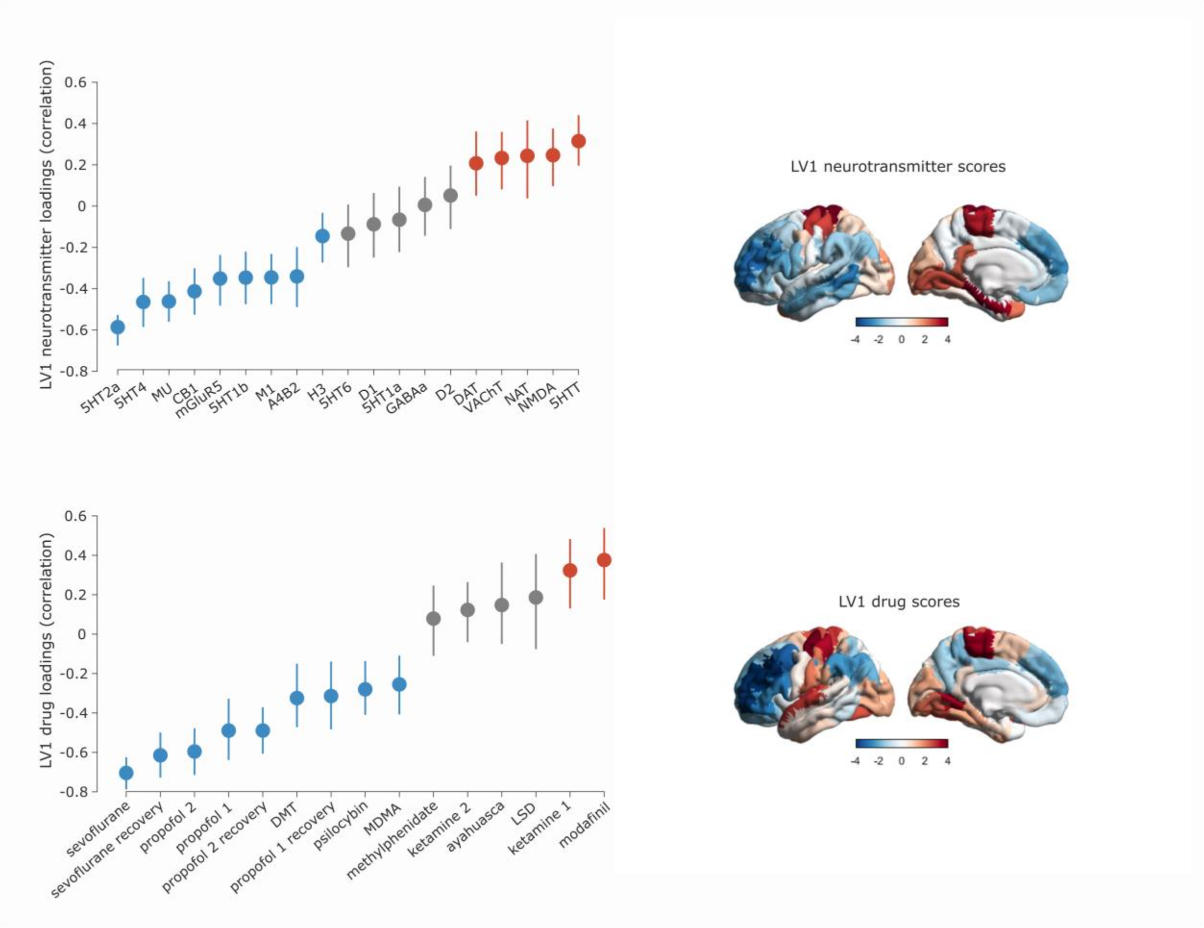


**Figure S5. Similar PLS1 is obtained with Lausanne-114 cortical parcellation.** Top: neurotransmitter loadings and scores. Bottom: drug loadings and scores.

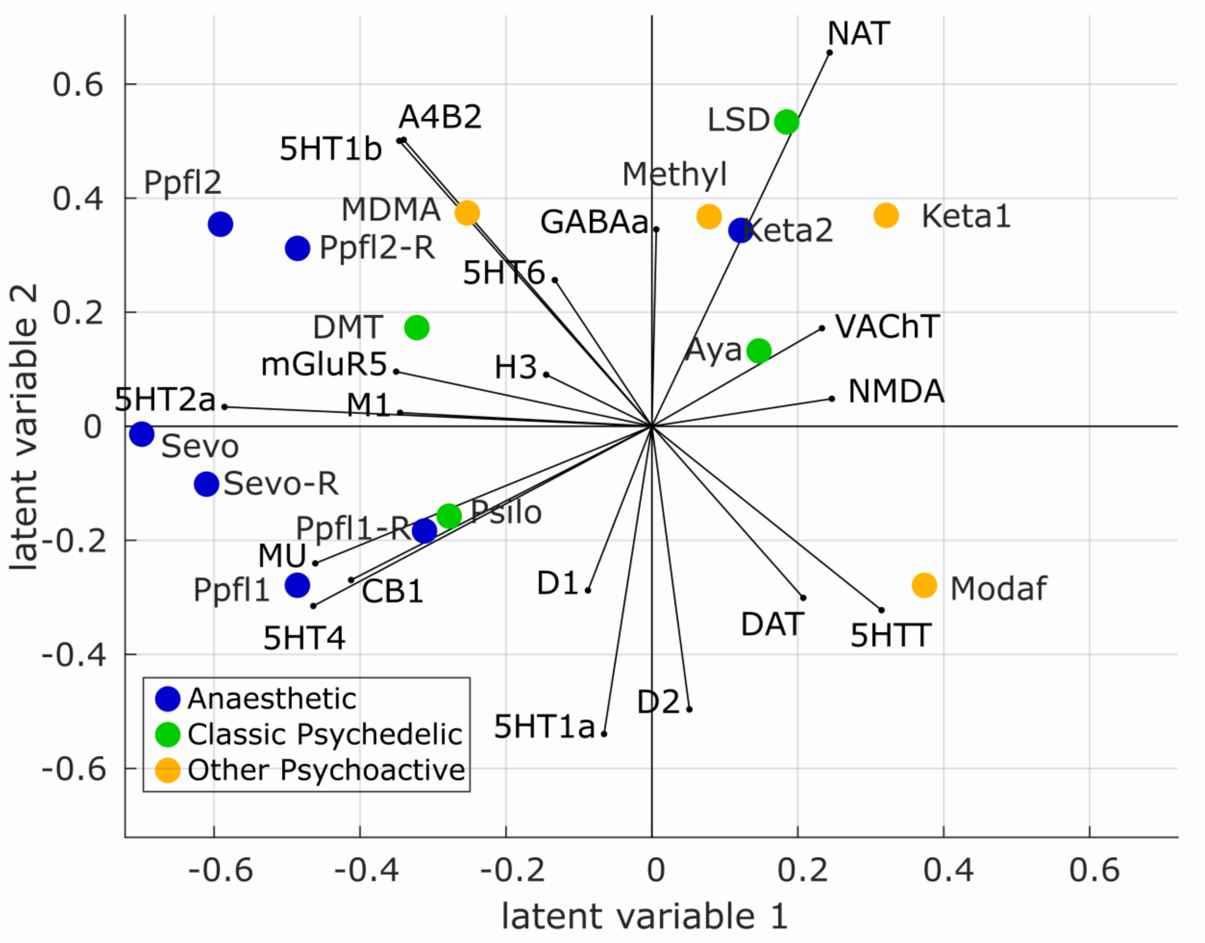


**Figure S6. Biplot for PLS1 and PLS2 obtained with Lausanne-114 cortical atlas.** Biplot shows neurotransmitters and pharmacological agents. Each drug is represented as a point reflecting its projection onto the first two latent variables of the PLS analysis, color-coded based on its effects on subjective experience (anaesthetic, classic psychedelic, or other psychoactive). Each neurotransmitter receptor and transporter is represented as a vector in the same 2D space.

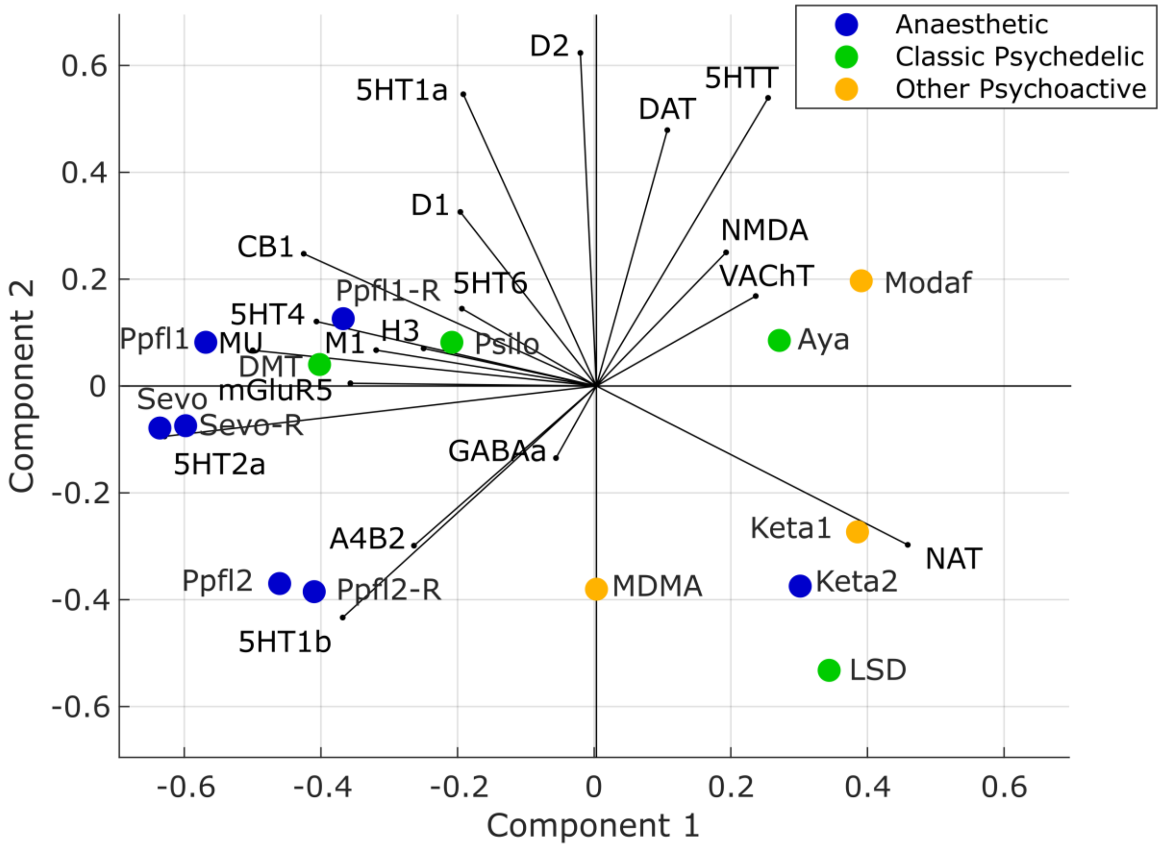


**Figure S7. Neurotransmitter landscape is qualitatively the same if methylphenidate data are not included.** Each drug is represented as a point reflecting its projection onto the first two latent variables of the PLS analysis, color-coded based on its effects on subjective experience (anaesthetic, classic psychedelic, or other psychoactive). Each neurotransmitter receptor and transporter is represented as a vector in the same 2D space.

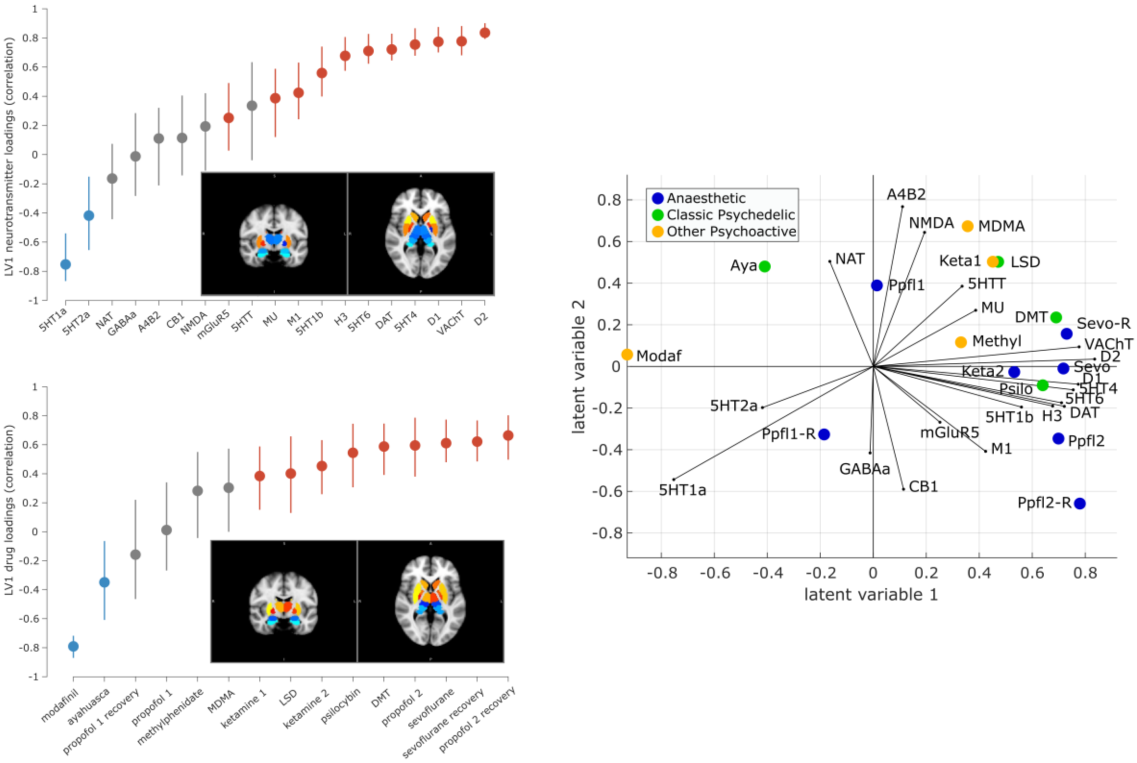


**Figure S8. Subcortical neurotransmitter-drug associations.** Left: neurotransmitter (top) and drug (bottom) loadings for PLS1, with corresponding PLS1 scores plotted on axial brain slices, for the subcortex (32 ROIs from the Melbourne atlas[177]). Right: biplot of neurotransmitters and pharmacological agents. Each drug is represented as a point reflecting its projection onto the first two latent variables of the PLS analysis, color-coded based on its effects on subjective experience (anaesthetic, psychedelic, or cognitive enhancer). Each neurotransmitter receptor and transporter is represented as a vector in the same 2D space.
